# Deciphering cytokine-driven ADP-ribosylation signaling networks via Af1521-based mass spectrometry analysis of labile Glu/Asp-linkages

**DOI:** 10.1101/2025.11.03.686237

**Authors:** Sara C. Buch-Larsen, Ivo A. Hendriks, Kyuto Tashiro, Jonas D. Elsborg, Sergey Y. Vakhrushev, Jesper V. Olsen, Bernhard Lüscher, Glen Liszczak, Ivan Ahel, Michael L. Nielsen

**Author notes:** Equal contribution (Sara C. Buch-Larsen, Ivo A. Hendriks).

## Abstract

ADP-ribosylation (ADPr) is a regulatory post-translational modification targeting nine amino acid residues. Glutamate- and aspartate-linked ADPr (Glu/Asp-ADPr) is chemically unstable during sample preparation for conventional mass spectrometry (MS)-based proteomics workflows, limiting its detection. Here, we systematically assessed the stability of ADPr linkages using synthetic peptides and confirmed that ester-linked Glu/Asp-ADPr is lost under alkaline conditions, elevated temperatures, or through hydrolysis by wildtype Af1521. We established an acidic enrichment workflow encompassing an Af1521 mutant, robustly preserving Glu/Asp-ADPr enabling their site-specific and systems-wide MS analysis. Applying this strategy to cytokine-stimulated A549 and HeLa cells, we identified >600 Glu/Asp-ADPr and >200 Cys-ADPr sites. Our analysis uncovered that Glu/Asp-ADPr marks distinct cytoplasmic protein networks enriched in immune functions, contrasting with Ser-ADPr typically observed on nuclear and chromatin-associated proteins. Quantitative profiling revealed reproducible ADPr patterns specific to cell type and treatment. Notably, PARP10 promoted Glu/Asp modification of ubiquitin, highlighting crosstalk between ADPr and ubiquitin signaling. Across interferon treatments, we identified a conserved network of antiviral PARPs and associated cofactors extensively modified on Glu/Asp residues, emphasizing residue-specific ADPr as a regulator of innate immune signaling. Together, our work establishes an MS-based proteomics workflow for identification of Glu/Asp-ADPr, provides a resource of site-specific modification events, and reveals residue-specific ADPr dynamics in immune signaling.

## INTRODUCTION

ADP-ribosylation (ADPr) is an emerging post-translational modification (PTM) that plays a crucial role in diverse cellular processes, including DNA repair, transcriptional regulation, and immune signaling (Luscher et al., 2018; Suskiewicz et al., 2023). It is catalyzed primarily by PARP enzymes (Luscher et al., 2022a), which transfer ADP-ribose moieties from NAD⁺ to the amino acid side chains of target proteins. Although ADPr was first described more than 60 years ago (Chambon et al., 1963), many aspects remain poorly understood and are still under debate – including fundamental questions such as which amino acids serve as predominant targets under specific cellular conditions.

Recent advances in mass spectrometry (MS)-based proteomics have transformed PTM research by enabling unbiased, proteome-wide analyses (Olsen and Mann, 2013). However, ADPr presents unique analytical challenges that have hindered its comprehensive characterization. It is highly dynamic and transient, typically of extremely low abundance, and structurally complex. ADPr can occur either as mono-ADPr, where a single ADP-ribose moiety is attached, or as poly-ADPr, in which ADPr chains are built upon pre-existing ADPr modifications (Luscher et al., 2018; Suskiewicz et al., 2023). Furthermore, it has been shown to modify at least nine different amino acid residues; Cys, Asp, Glu, His, Lys, Arg, Ser, Thr, and Tyr (Larsen et al., 2018). We have previously demonstrated that ADPr is exceptionally labile under higher-energy collisional dissociation (HCD) fragmentation, and thus for accurate site localization the non-ergodic fragmentation propensity of electron-transfer dissociation (ETD)-based approaches is required (Larsen et al., 2018).

Over the past decades, several MS-based strategies have been developed to tackle these challenges, each with distinct advantages and limitations (Bilan et al., 2017; Daniels et al., 2014; Gagné et al., 2015; Hendriks et al., 2019; Martello et al., 2016; Zhang et al., 2013). Some methods offer broad coverage but reduced specificity, while others achieve high confidence in site localization at the cost of sensitivity or enrichment efficiency. Consequently, these complementary approaches have collectively advanced the field but often capture only partially overlapping subsets of the ADPr landscape. A major breakthrough came with the identification of serine ADPr, particularly in the DNA damage response, mediated by PARP1 and PARP2 in concert with the cofactor HPF1 (Bonfiglio et al., 2017; Fontana et al., 2017; Palazzo et al., 2018; Suskiewicz et al., 2020). Using Af1521 macrodomain-based enrichment strategies, we and others have developed MS workflows that robustly detect serine-linked ADPr, greatly expanding knowledge of its regulatory roles (Bilan et al., 2017; Hendriks et al., 2019; Larsen et al., 2018). Despite this progress, other physiologically relevant ADPr linkages remain underexplored. In particular, glutamate- and aspartate-linked ADPr have proven especially challenging to study. It has recently become clear that these acidic residue linkages are highly labile under the mildly basic conditions (pH 8-8.5) routinely used in standard MS-based workflows for protein lysis and digestion (Tashiro et al., 2023). This property likely explains their systematic underrepresentation in previous datasets and underscores the need for tailored analytical strategies to study these modifications.

Here, we present an MS-based workflow designed specifically for the detection of Glu- and Asp-linked ADPr. This strategy overcomes key technical limitations and enables confident site-specific identification of these modifications. Applying this approach, we performed in-depth and system-wide screens of Glu/Asp-linked ADPr in response to cytokine treatment, in both A549 cells and HeLa cells with induced PARP10 expression, uncovering insights into ADPr regulation in immune signaling and provide a rich resource for investigating residue-specific ADPr linkages in diverse biological contexts.

## RESULTS

### ADPr linkages differ markedly in stability across proteomics sample preparation conditions

Recently, the lability of ester-linked ADP-ribosylation was demonstrated in the context of alkaline or high-temperature conditions (Javed et al., 2023; Longarini and Matic, 2024; Tashiro et al., 2023). Over the last decade, we and others have performed multiple mass spectrometry (MS) screens with the routine usage of alkaline conditions and long incubations at room temperature or above during sample preparation (Bilan et al., 2017; Buch-Larsen et al., 2020; Hendriks et al., 2019; Larsen et al., 2018; Zhang et al., 2013), suggesting that ester-linked ADPr (i.e. Glu/Asp-ADPr) has been poorly conserved. Here, we set out to systematically assess the stability of ADPr linkages to four different amino acid residues, in the context of standard MS-based proteomics sample preparation conditions and buffers (Fig. 1A). For this purpose, we used synthetic peptides with a largely homologous sequence, with ADPr linked specifically to an Asp, Glu, Arg, or Ser residue. The peptide sequences were non-isobaric to allow distinguishing them easily even from mixtures, and all analysis was performed with the ETD-capable Lumos mass spectrometer to allow confident identification and localization of the ADPr within each peptide.

**Figure 1.**
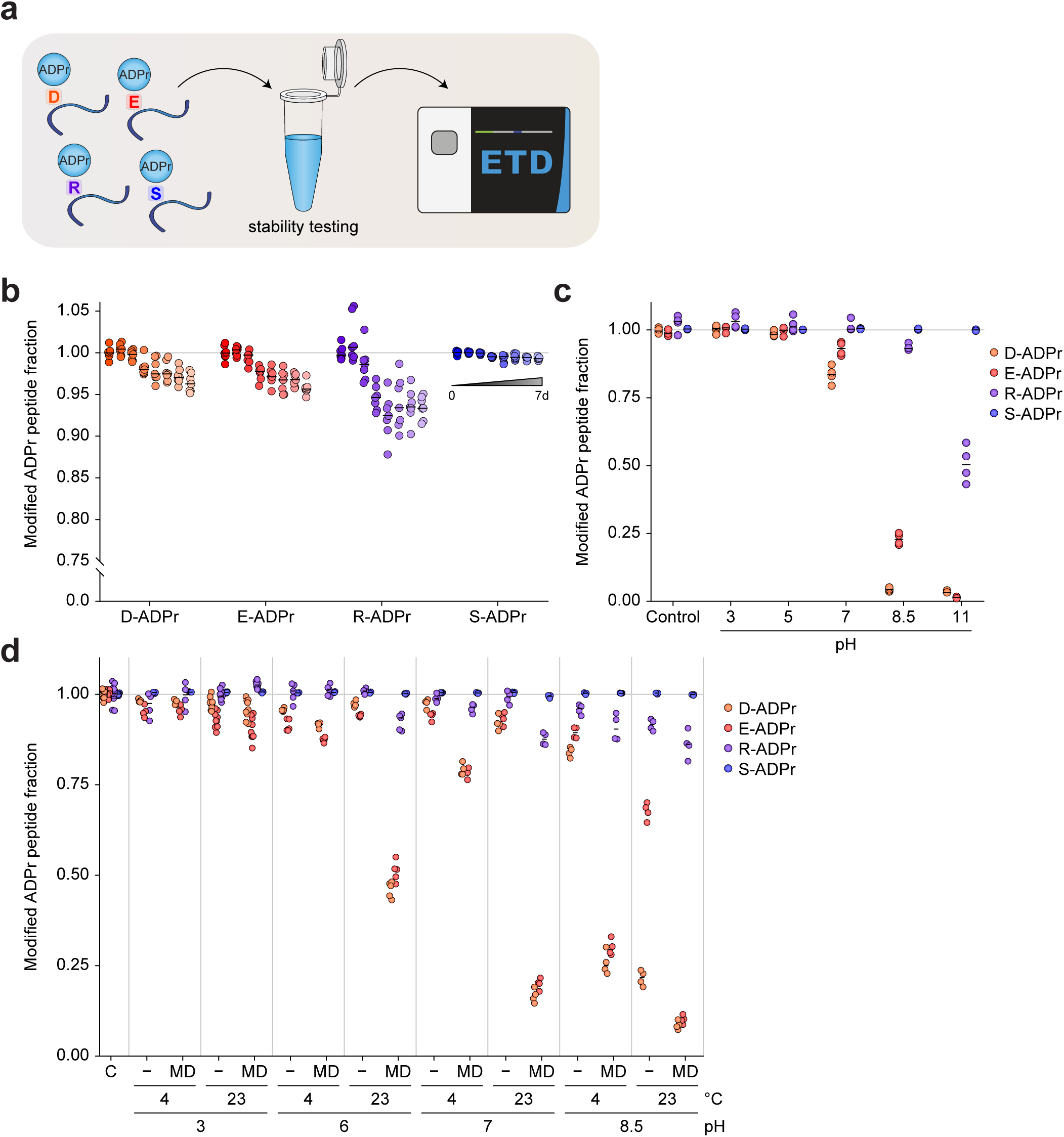
MS-based evaluation of stability of ADPr linked to Asp, Glu, Arg, and Ser residues. (A) Summary of the strategy employed. Synthetic peptides bearing specific APDr linkages were incubated in a variety of ways, prior to analysis using ETD-based mass spectrometry. (B) Relative stability of ADPr linkages over time at 7 °C, normalized to timepoint 0. Median values are indicated, *n*=7 technical replicates. (C) Relative ADPr stability across different pH values across common MS buffers at room temperature, prior to vacuum drying and analysis, *n*=4 technical replicates. (D) Relative ADPr stability at different pH, temperature, and in the absence of presence of Af1521 macrodomain. *n*=4 technical replicates.

We first investigated the overall stability of the four different linkages over time, by incubating them for up to one week in a high-performance liquid chromatography (HPLC) system under standard proteomic sample conditions, i.e. in 0.1% formic acid in a 96-well plate kept at 7 °C (Fig. 1B). Reassuringly, we found that all four linkages are highly stable under these conditions, with at most ∼5% degradation for Arg-ADPr over the course of one week, supporting that LC-MS conditions are not problematic for the analysis of any ADPr linkage type. Next, we incubated the peptide mixtures for three hours in buffers with varying pH (Fig. 1C), and confirmed the previously observed lability of Glu/Asp-ADPr at alkaline conditions. Notably, even under physiological conditions ester-linked ADPr is not entirely stable, and furthermore we also observed that half of Arg-ADPr was reversed under highly alkaline conditions that are typically employed during high-pH fractionation (Hendriks et al., 2019). Acidifying the mixtures prior to vacuum concentrating them at 60 °C was only able to partially prevent hydrolysis of ADPr (Fig. S1A). As a control, we also incubated the ADPr peptide mixture using hydroxylamine, either pH-neutralized or not (Fig. S1B). Indeed, hydroxylamine very rapidly reversed Glu/Asp-ADPr, whereas Ser-ADPr was unaffected. Alarmingly, we found that Arg-ADPr was also affected by hydroxylamine, with ∼25% loss after 15 min, ∼50% loss after 3 h, and essentially total loss after 24 h. Finally, we combined the ADPr peptide mixture with the routinely applied Af1521 macrodomain (Fig. 1D), which has been reported to exhibit Glu/Asp-ADPr hydrolase activity (Jankevicius et al., 2013; Rosenthal et al., 2013). We observed that at neutral pH and at room temperature, Af1521 was able to largely abolish Glu/Asp-ADPr, and even at 4 °C was able to slightly reverse these linkages. At pH 8.5, commonly used during proteomics sample preparation, Af1521 can efficiently reverse Glu/Asp-ADPr even at 4 °C. At pH 6, Af1521 could still affect ester-linked ADPr at room temperature, but the mixture was stabilized at 4 °C. Taken together, these findings highlight the pronounced sensitivity of Glu/Asp-linked ADPr to alkaline conditions, underscoring the need for carefully controlled sample preparation to preserve all ADPr linkage types.

### Acidic workflow and mutant Af1521 uncover hidden Glu/Asp-ADPr

Based on our observations using the synthetic ADPr peptides, we set forth to adapt and benchmark our Af1521 enrichment methodology, aiming to establish a MS-based strategy that can stably purify and identify the labile ester-linked Glu/Asp-ADPr (Fig. 2A). To this end, we utilized three distinct and relevant cellular systems that putatively encompass a wide range of different ADPr linkage types. These included interferon-treated A549 cells; where MAR is expected through PARP14 activation (Kar et al., 2024; Ribeiro et al., 2024), H_2_O_2_-treated HeLa cells; where a large accumulation of Ser-ADPr is expected (Larsen et al., 2018; Leidecker et al., 2016; Liu et al., 2017; Palazzo et al., 2018); and H_2_O_2_-treated U2OS HPF1 knockout cells; wherein a shift towards Glu/Asp-ADPr is anticipated (Langelier et al., 2021; Palazzo et al., 2018; Rudolph et al., 2021). We either performed our established Af1521-based workflow (Hendriks et al., 2019), which relies on alkaline conditions, or otherwise we utilized our modified Af1521 workflow under acidic conditions (pH <6.3) at all stages while also reducing incubation times and temperatures to a minimum. Furthermore, we evaluated both approaches in the absence or presence of PARG, and utilized either wildtype or mutant (K35E / Y145R) Af1521 macrodomain (Garcia-Saura et al., 2021; Nowak et al., 2020)(Fig. 2B).

**Figure 2.**
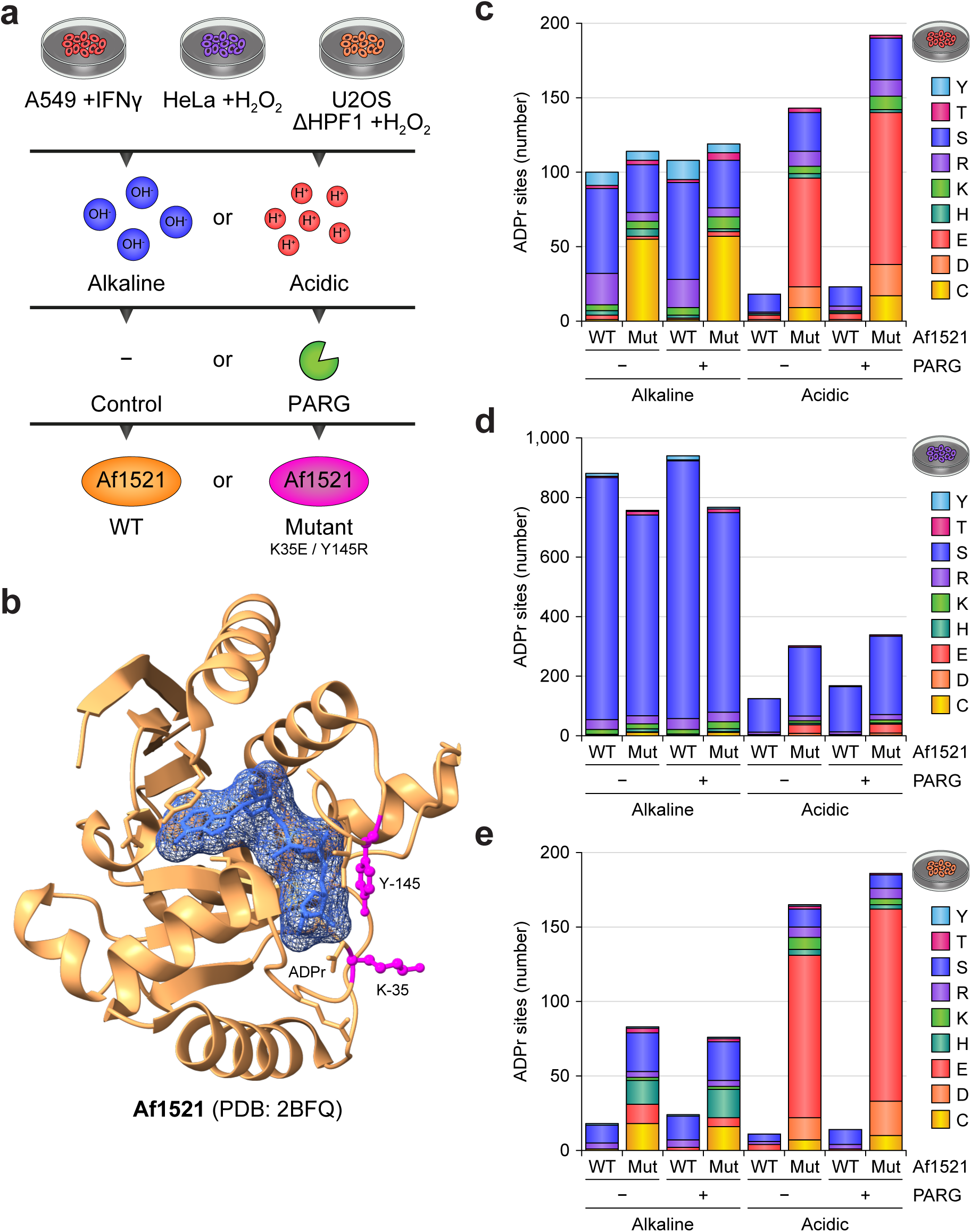
Benchmarking different proteomics workflows for purification of any ADPr linkage type. (A) Experimental design overview. Different cell lines were treated as indicated, prior to lysis and further processing under either predominantly alkaline conditions, or exclusively acidic conditions. Following proteolytic digestion, peptides with treated with PARG or not, and enrichment of ADPr was performed using either wildtype of mutant Af1521 macrodomain. (B) Structure of the Af1521 macrodomain, as reproduced from PDB structure 2BFQ. The location of a bound ADPr moiety, as well as the Lys-35 and Tyr-145 residues that are mutated in the engineered macrodomain (Garcia-Saura et al., 2021; Nowak et al., 2020), are indicated. (C) Overview of the cumulative number of ADPr sites identified in the A549 +IFNγ samples, with distribution of amino acid types indicated. (D) As **C**, but for HeLa +H_2_O_2_ samples. (E) As **C**, but for U2OS ΔHPF1 +H_2_O_2_ samples.

All samples were analyzed using ETD-based mass spectrometry and data was processed in an unrestricted manner where ADPr is theoretically allowed to reside on any of the twenty naturally occurring amino acid residues. Overall, we found that both the alkaline and acidic methods were comparably efficient at generating samples that facilitated MS-based localization of the ADPr moiety to specific amino acid residues (Fig. S2A), with approximately two-thirds of peptide-spectrum-matches (PSMs) localized at a >90% probability. Furthermore, across all samples investigated, we were able to confidently detect (>99% localization) ADPr on the nine previously reported reactive amino acids (Larsen et al., 2018), with no strong evidence for modification of any further residue types (Fig. S2B). Thus, all further data analyses were performed with ADPr allowed to reside on Cys, Asp, Glu, His, Lys, Arg, Ser, Thr, and Tyr residues.

In the context of interferon-treated A549 cells, we were able to identify several hundreds of ADPr sites (Table S1). Intriguingly, the distribution of linkage types varied dramatically depending on the purification strategy used, with Glu/Asp-ADPr exclusively detected using the acidic workflow (Fig. 2C and S2C), with Glu as the main acceptor site. Strikingly, this was only observed when using the mutant Af1521, suggesting this variant of the macrodomain could bind stronger and/or hydrolyzes Glu/Asp-ADPr with a lower efficiency. The combination of wildtype Af1521 and acidic conditions resulted in mainly unmodified peptides being detected (Fig. S2D), with ADPr peptide purity at ∼10% otherwise. Notably, Cys-ADPr was mainly observed when using the mutant Af1521, and prominently under alkaline conditions.

When considering H_2_O_2_-treated HeLa cells, we could readily reproduce the preponderant presence of Ser-ADPr, with just under 1,000 sites detected (Fig. 2D and S2E and Table S1). Regardless of the purification workflow, Ser-ADPr remained dominant especially by abundance, although interestingly the efficiency of Ser-ADPr purification was greatly diminished from the alkaline (∼80% purity) to the acidic (∼5-25% purity) workflow (Fig. S2F). The mutant Af1521 was overall able to increase the total abundance of purified ADPr (Fig. S2E), which paradoxically decreased the number of identified sites under alkaline conditions while increasing the number under acidic conditions (Fig. 2D).

In the absence of HPF1, the transfer of ADPr to Ser is greatly diminished (Bonfiglio et al., 2017; Hendriks et al., 2021; Palazzo et al., 2018). Indeed, HPF1 knockout cells in response to H_2_O_2_ treatment, we observed a dramatic reduction in the number of Ser-ADPr sites and their overall abundance (Fig. 2E and S2G and Table S1), compared to a HPF1-proficient system (Fig. 2D and S2E). Utilizing the mutant Af1521 and the alkaline workflow revealed the presence of additional ADPr, mostly targeting Cys and His residues. Moreover, the combination of the mutant Af1521 and the acidic workflow uncovered a considerable number of Glu/Asp-ADPr sites, representing ∼90% of the overall ADP-ribosylome, hereby demonstrating a molecular switch from Ser to Asp/Glu targeting. Interestingly, across all three cellular systems, we did not observe any notable effect of incubating the samples with PARG (Fig. 2C-E). Further, we noted that in all cellular contexts and while using the mutant Af1521, we could achieve a peptide purity of at least 10-20%, which is on par or superior to antibody-based enrichments (Hendriks et al., 2021).

Overall, we demonstrate that our strategy utilizing the Af1521 macrodomain and either alkaline or acidic conditions, can capture the heterogeneity of ADPr acceptor sites entailed within diverse biological systems.

### Ser and Glu/Asp-ADPr target distinct biological and subcellular proteins

Across all conditions we investigated during the benchmark experiments (Fig. 2), we observed ADPr to most frequently target Ser, Glu/Asp, Arg, and Cys residues (Fig. 3A). Although the comparison was technical in nature with a focus on different lysis and purification conditions, and was performed from one sample batch, we nevertheless explored how the different ADPr linkage types correlated to their respective biological origins.

**Figure 3.**
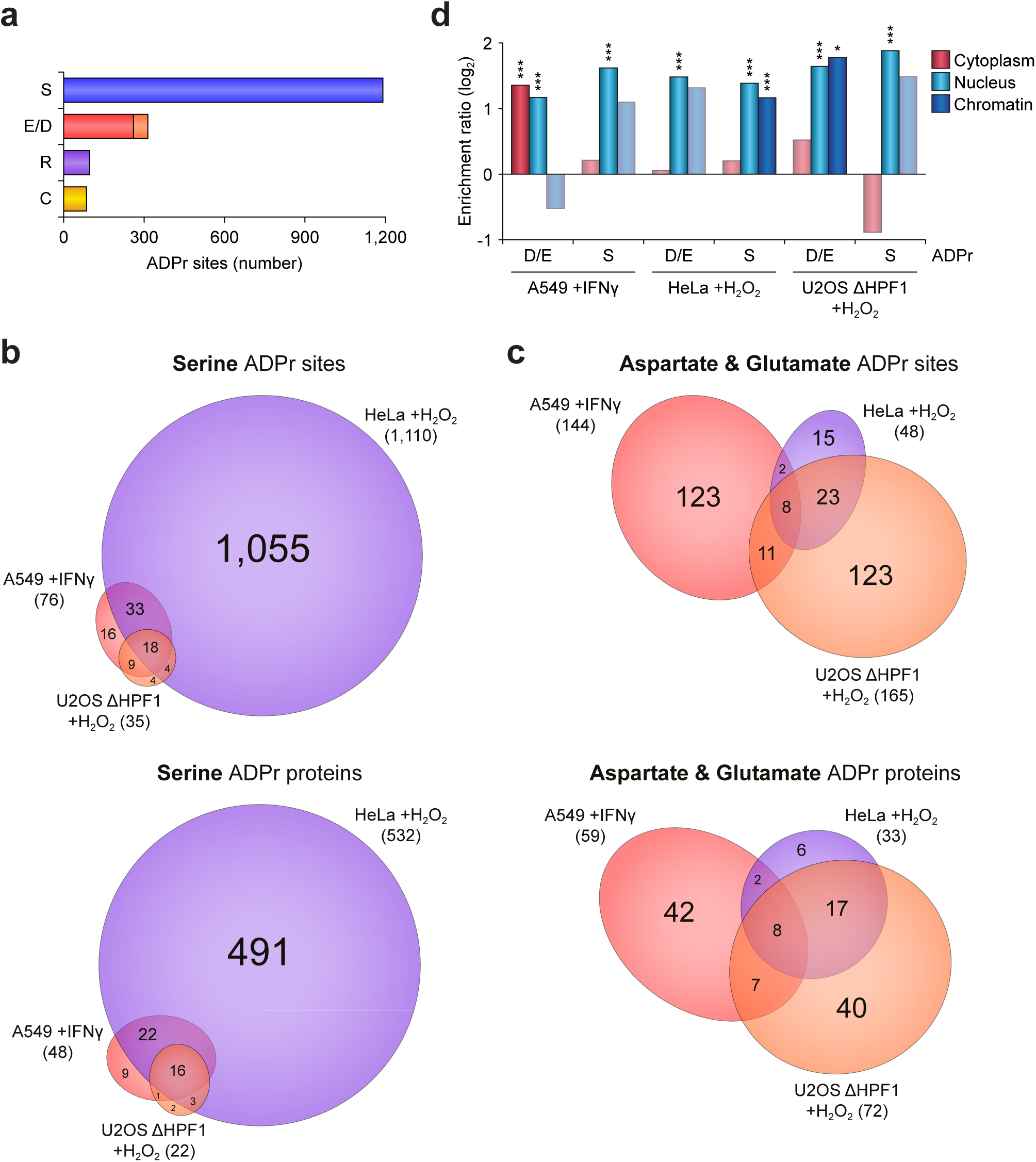
Distribution of different ADPr linkages across different cell lines and treatments. (A) Total number of ADPr sites identified in the benchmark experiment, for the five most observed linkages. (B) Scaled Venn diagram showing overlap between the different cell types and treatments, for either Ser ADPr sites (top) or Ser ADPr target proteins (bottom). (C) As **B**, but for Asp and Glu ADPr. (D) Term enrichment analysis, where the Gene Ontology subcellular localization was compared for subsets of ADPr target proteins, versus the background total proteome. Positive enrichment ratio indicates a higher-than-expected observation of ADPr target proteins within the category. * *p*<0.05, *** *p*<0.0001, determined via two-tailed Fisher Exact testing with Benjamini-Hochberg correction for multiple-hypotheses testing. Faded bars lack statistical significance.

As anticipated, considering the cumulative number over all conditions, ADPr on serine residues was the most frequently observed (Fig. 3A), with the vast majority found in H_2_O_2_-treated HeLa cells (Fig. 3B). In the absence of HPF1, or in A549 cells that were not treated with H_2_O_2_, Ser-ADPr only targeted a few proteins. In these cases, there was still a considerable overlap to Ser-ADPr detected in H_2_O_2_-treated HeLa. In case of Glu/Asp-ADPr, in contrast to Ser-ADPr, the majority of sites and target proteins were observed in IFN-treated A549 and H_2_O_2_-treated U2OS lacking HPF1 (Fig. 3C). Interestingly, despite similar numbers of Glu/Asp-ADPr sites identified in both of these systems, their overlap was relatively small at both the site and the protein level, hinting at distinct biological pathways being targeted. For Cys-ADPr, most detections occurred in IFN-treated A549, with only a few in the other two systems (Fig. S3A). Interestingly, whereas for Glu/Asp-ADPr we found on average 3 sites-per-protein, for Cys-ADPr the majority of target proteins contained exactly one site. We noted Arg-ADPr to occur somewhat sporadically across the three cell lines, with an overall poor overlap (Fig. S3B), which may be due to different expression levels of ARTC enzymes (Glowacki et al., 2002).

For the two largest groups of ADPr sites, Ser and Glu/Asp, we investigated the annotated subcellular localization of their respective target proteins (Fig. 3D). As expected, we found that Ser-ADPr significantly targeted nuclear proteins across all biological systems investigated, consistent with writers of this modification localizing to the nucleus (Gibbs-Seymour et al., 2016). In case of H_2_O_2_ treatment in HPF1-proficient cells, chromatin-associated proteins were also a significant target. Intriguingly, in HPF1-deficient cells, the chromatin targeting significantly shifted to Glu/Asp-ADPr, suggesting that the same proteins or biological pathways may remain targeted by ADPr, but with altered amino acid specificity. Finally, considering IFN-treated A549, the Glu/Asp-ADPr uniquely and significantly targeted cytoplasmic proteins, demonstrating that certain linkage types can target different subcellular pathways or compartments depending on biological context.

### Quantitative profiling reveals robust and reproducible Glu/Asp-ADPr landscapes

Having established a strategy able to judiciously identify glutamate- and aspartate-linked ADPr, we wanted to validate our approach in a quantitative manner. To this end, we set up a large-scale quadruplicate experiment wherein we compared untreated A549 cells with IFNα- or IFNγ-treated cells. Furthermore, we included HeLa cells which allow for selective expression of PARP10, which is a prominent Glu/Asp mono-ADPr transferase (Garcia-Saura and Schuler, 2021; Kaufmann et al., 2015), and performed either mock, IFNα, or TNFα treatment (Fig. 4A). For enrichment of ADPr, we utilized our acidic workflow employing mutant Af1521, and analyzed all samples using ETD-based mass spectrometry.

**Figure 4.**
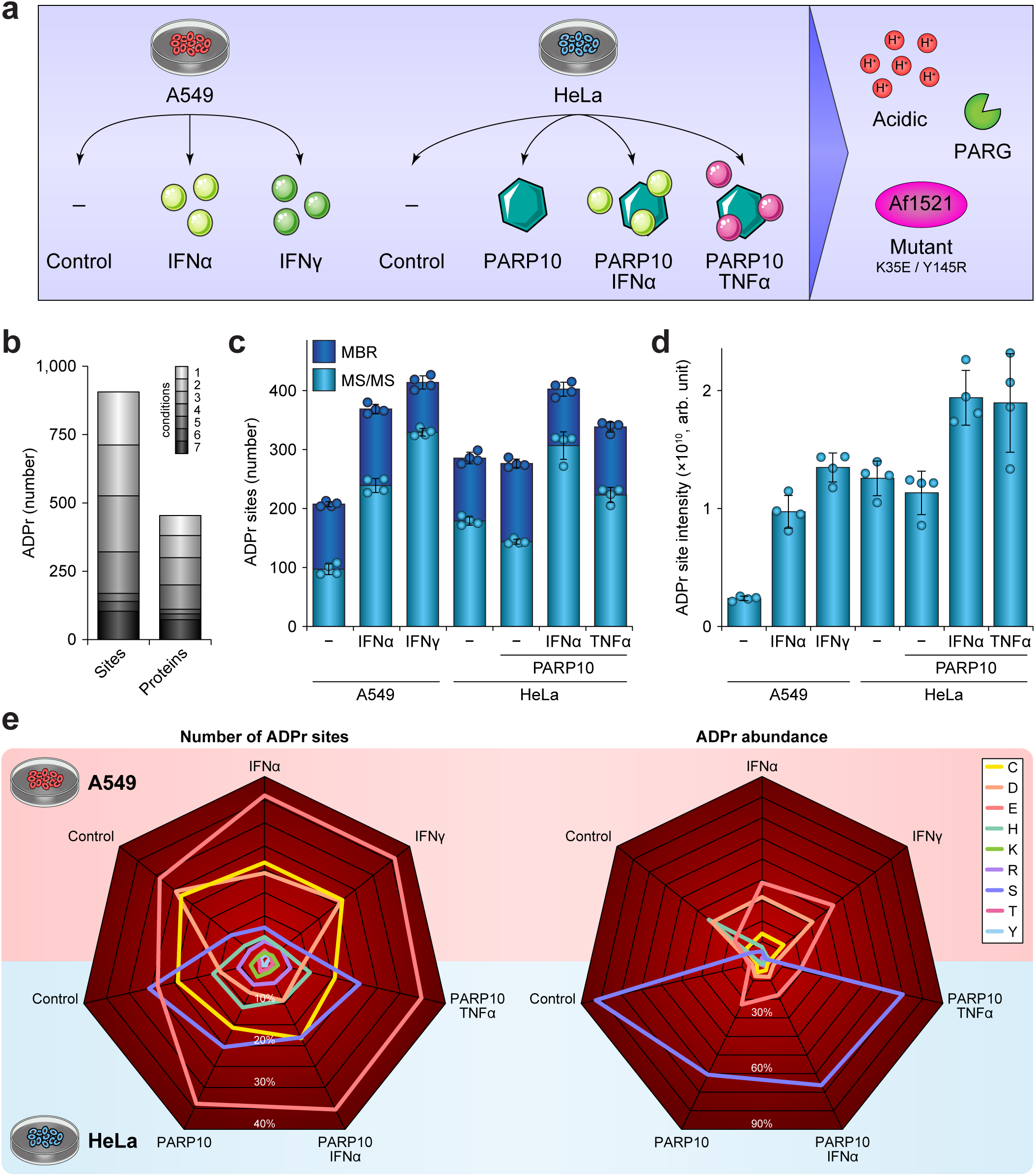
Quantitative profiling of labile ADPr sites in the context of cytokine treatment. (A) Overview of the experimental design. Briefly, A549 and HeLa cells were cultured in quadruplicates, and treated as indicated. Lysis and further processing was performed under exclusively acidic conditions, with enrichment using mutant Af1521, to preserve labile ADPr linkages. Analysis was performed using ETD-based mass spectrometry, *n*=4 cell culture replicates. (B) Overview of the total number of ADPr sites and target proteins identified across the quantitative experiment. (C) Number of ADPr sites identified per experimental condition, either directly within each sample (MS/MS), or after matching of MS-based evidence between other replicates and conditions (MBR). Error bars represent s.d., *n*=4 cell culture replicates. (D) As **C**, but visualizing total abundance of ADPr site signal. (E) ‘Spiderman’ plot, visualizing the relative distribution of each residue-specific ADPr to the total ADPr within the experiment, either when considering the number of sites (left plot), or when going by summed abundance (right plot). A549 conditions are visualized at the top, and HeLa conditions at the bottom.

In total, we identified 906 high-confidence ADPr sites residing on 454 target proteins (Fig. 4B, Table S2, and Table S3), with approximately two-thirds of proteins modified by a single ADPr (Fig. S4). Only a small subset of ADPr sites was found in all conditions, highlighting the biological variability between the conditions (Fig. 4B). We reproducibly identified between 200 and 400 ADPr sites per condition, with the higher numbers observed after cytokine treatment (Fig. 4C), although we observed a higher baseline ADPr abundance in HeLa cells (Fig. 4D). Overall, ADPr most frequently targeted Glu, Asp, Cys, and Ser residues, with the latter responsible for the higher baseline in HeLa compared to A549 cells (Fig. 4E). Consistent with the targeting of ADPr to Glu/Asp in our benchmark experiment (Figs. 2-3), we found that modification in A549 cells primarily targeted Glu/Asp, especially following interferon treatment. When expressing PARP10 in HeLa cells, Glu-ADPr was prominently induced, in lieu of the Ser-ADPr that was otherwise notably present in HeLa cells. In all cases, Cys-ADPr represented ∼20% of all sites, and ∼5-10% of overall ADPr abundance. Taken together, our data reveal highly distinct patterns in which amino acid types are targeted by ADPr, with either widespread modification of numerous residues or focused modification of a few sites, depending not only on treatment condition, but moreover on the cellular system employed.

We observed an overall high degree of reproducibility, with average Pearson correlations of 0.91 to 0.96 within same-condition replicates (Fig. S5). As expected, correlation was lower across different treatment conditions within the same cell type, and much lower across different cell lines. Hierarchical clustering of the data further illustrated a high degree of same-condition replicate reproducibility (Fig. 5A), while visualizing distinct clusters of ADPr target proteins. Principle component analysis revealed that the primary degree of variance originates from the different cell lines, followed by untreated versus cytokine-treated, with only modest variance within treatment clusters (Fig. 5B).

**Figure 5.**
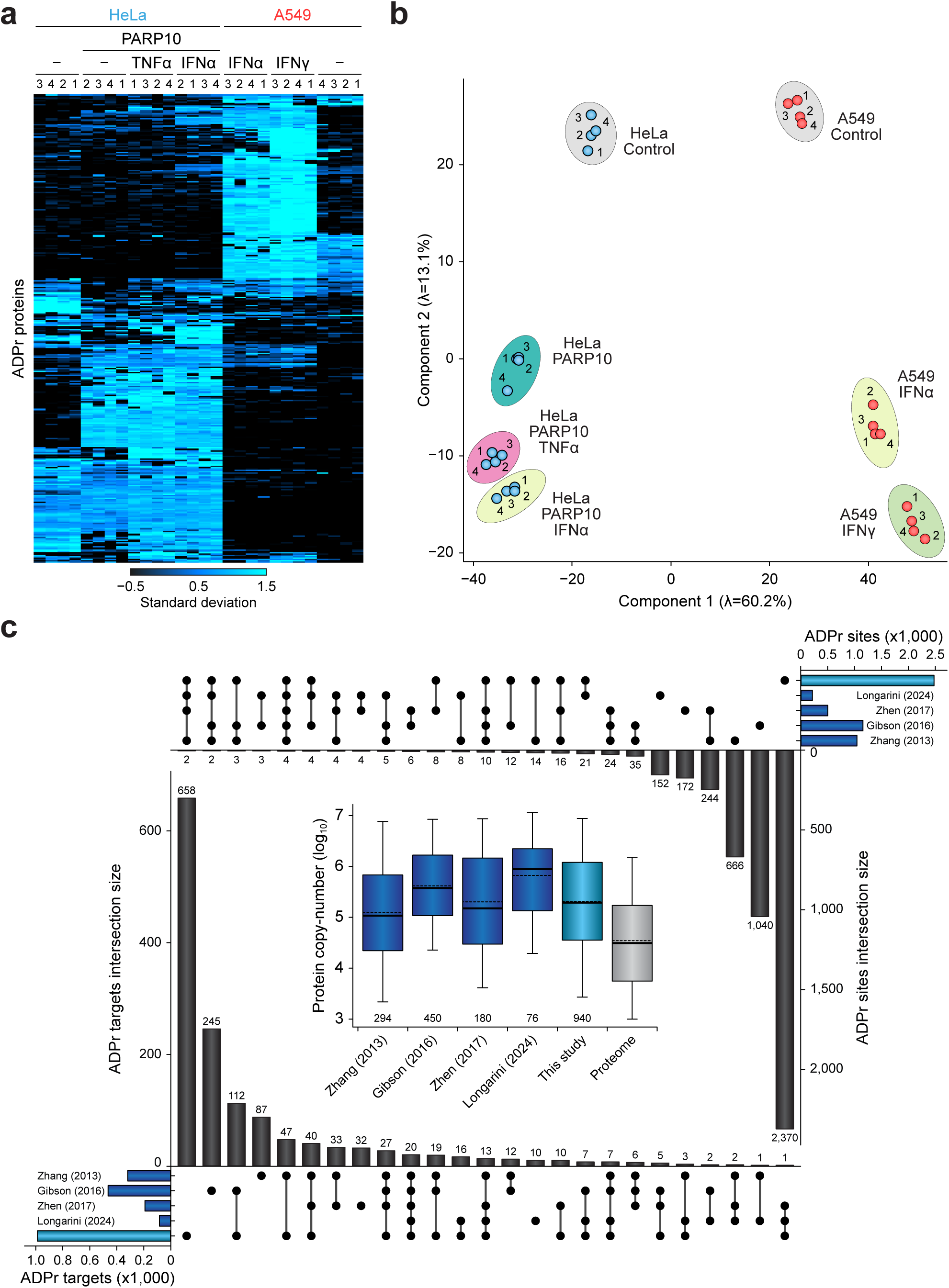
Reproducibility analysis and comparing to previous Glu/Asp ADPr studies. (A) Unsupervised hierarchical clustering analysis, resulting in a heatmap which displays the overall presence or absence of ADPr target proteins as identified across all replicates and conditions. Summed ADPr site abundances were z-score normalized prior to Euclidian clustering. (B) Principle component analysis, visualizing the greatest degrees of variance (component 1 followed by component 2) within the entire dataset. (C) UpSet analysis for ADPr sites (top-right), UpSet analysis for ADPr target proteins (bottom-left), and depth of sequencing analysis (central). The UpSet analyses demonstrate which studies, or combinations of studies, (co-)identified the most ADPr sites or proteins, with the top 25 most occurring combinations visualized. The depth of sequencing is a boxplot visualization of the iBAQ protein expression values, derived from Bekker-Jensen et al. (Bekker-Jensen et al., 2017), for all ADPr target proteins identified in each study. Thick line; median, dashed line; mean, box limits; 1^st^ and 3^rd^ quartile, whisker limits; 5^th^ and 95^th^ percentile. The number below the boxes denotes the number of ADPr target proteins within the study. Previously published ADPr data were adapted from Zhang et al. (Zhang et al., 2013), Gibson et al. (Gibson et al., 2016), Zhen et al. (Zhen et al., 2017), and Longarini and Matic (Longarini and Matic, 2024).

Since we developed a methodology capable of preserving labile Glu/Asp-linked ADPr, and profiled several hundreds of Glu/Asp-ADPr sites across both our benchmarking and quantitative experiment, we wanted to evaluate the comprehensiveness of our profiling. To this end, we compared all ADPr sites mapped in this work (Fig. 5C and Table S4), to previous MS-based proteomics studies that facilitated Glu/Asp-ADPr identification (Gibson et al., 2016; Longarini and Matic, 2024; Zhang et al., 2013; Zhen et al., 2017). Three of these studies relied on hydroxylamine-derivatization of Glu/Asp-ADPr prior to MS using collisional fragmentation (Gibson et al., 2016; Zhang et al., 2013; Zhen et al., 2017), whereas the most recent study similarly to us relied on electron-based fragmentation of peptide bearing the intact ADPr moiety (Longarini and Matic, 2024). We first assessed the depth of sequencing achieved across these studies via comparison of known protein copy-numbers and correlating these to the identified ADPr target proteins (Fig. 5C). Overall, our experiments reached a similar depth of sequencing compared to the two studies from the Yu lab, and achieved greater depth than the other ADPr studies in the comparison (Table S4). When considering overlaps in ADPr target proteins across studies, whereas there were some large groups of study-specific proteins, we overall see a very strong overlap between our work and the three studies with most ADPr identifications. Interestingly, at the site-specific level, our study overlaps best with the most recent, smaller study, which also relied on an acidic sample preparation workflow and ETD-based fragmentation (Fig. 5C). Taken together, we identified >600 high-confidence Glu/Asp-linked ADPr sites across conditions and cell types with strong reproducibility (Tables S1 and S2). Results showed cell line- and treatment-specific ADPr patterns, with robust protein-level overlap to previous studies, confirming both the depth and reliability of our profiling approach.

### PARP10 rewires the ADP-Ribosylome and targets ubiquitin for ADPr

Expression of PARP10 in HeLa cells did not result in a difference in the overall number of ADPr sites identified, nor did the overall ADPr abundance change (Fig. 4C-D). However, based on the hierarchical clustering and principal component analysis, there was a considerable and consistent change in the overall ADP-ribosylome following PARP10 induction (Fig. 5A-B). Indeed, volcano plot analysis demonstrated a significant re-wiring, with 61 proteins upregulated and 47 proteins downregulated in their ADP-ribosylation (Fig. 6A). Overall, 47 (∼77%) of the upregulated proteins were annotated as cytosolic, whereas 42 (∼89%) of the downregulated proteins were annotated as nuclear (Table S3). Reassuringly, PARP10 itself was found to be ADP-ribosylated upon expression. Further treatment of this cellular system with either IFNα or TNFα resulted in significant upregulation of 55 or 13 proteins (Fig. 6B and S6), respectively, with only five proteins downregulated in case of IFNα treatment. Altogether, while treatment with IFNα induced a robust effect with modification of the PARP9/PARP14/DTX3L axis (Dukic et al., 2023; Kar et al., 2024; Ribeiro et al., 2024), TNFα did not elicit a notable response within our experimental setup.

**Figure 6.**
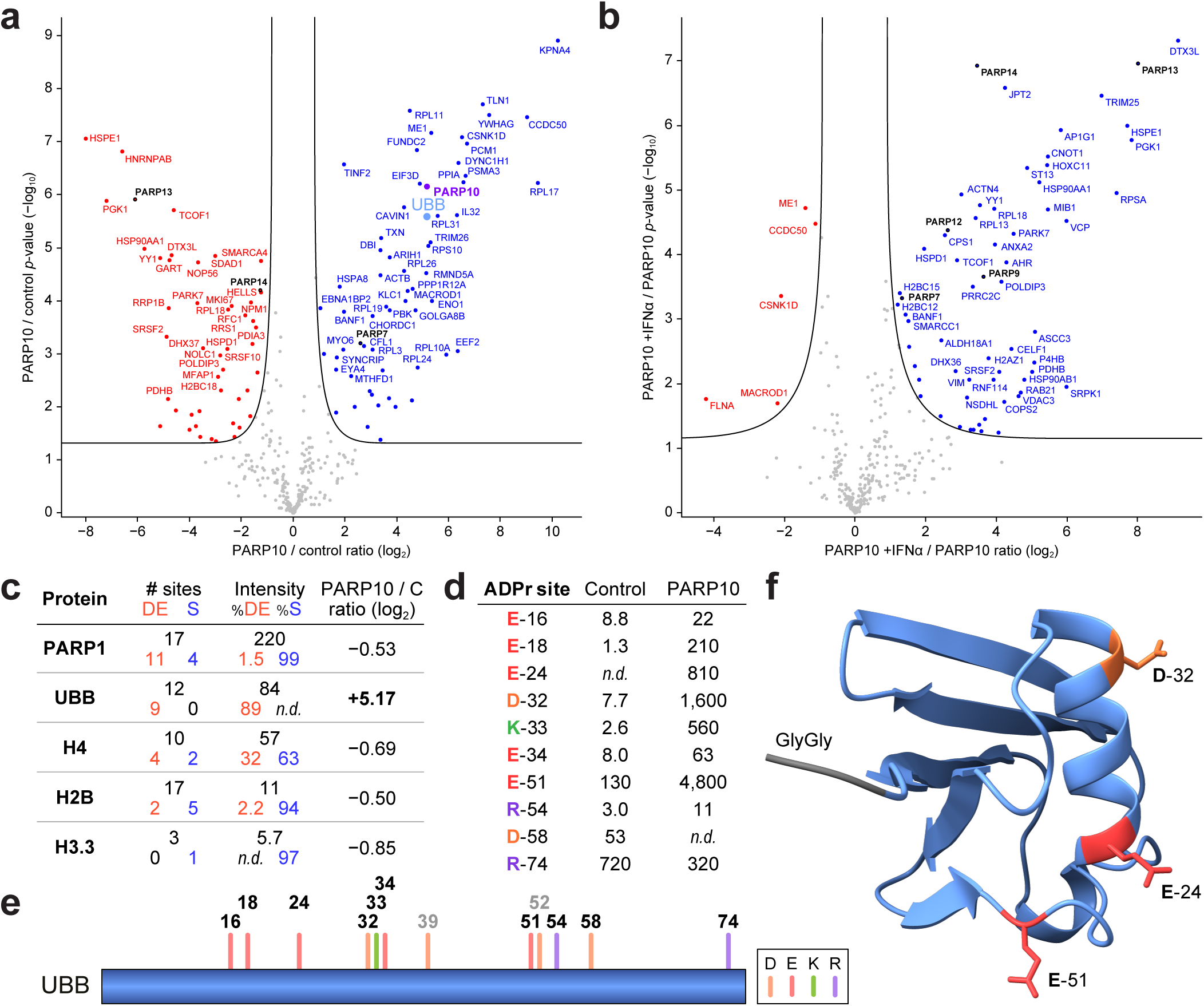
PARP10 expression induces ubiquitin ADP-ribosylation. (A) Volcano plot analysis, visualizing ADPr target protein dynamics between PARP10-induced (right) and control (left). The significance cut-off line is drawn at *q*=0.01. Significance determined via student’s two-sided t-testing with permutation-based FDR control, *n*=4 cell culture replicates. (B) As **A**, but for PARP10-induced +IFNα (right) versus PARP10 (left). (C) Overview of the top five most abundantly ADPr proteins in HeLa cells, following PARP10 induction. Number and abundance (in arbitrary units) of Glu/Asp and Ser ADPr, and overall ratio of change in modification, is indicated for each of the proteins. (D) Overview of all ADPr sites detected on ubiquitin, and their abundance (in arbitrary units), in control and PARP10-induced cells. (E) Linear visualization of ubiquitin with ADPr sites highlighted. (F) Structure of ubiquitin, with the three ADPr sites most prominently induced on PARP10 induction highlighted.

PARP10 expression resulted in a considerable shift in the ADP-ribosylome, and within this context we investigated which proteins overall were the most highly modified by ADP-ribosylation (Fig. 6C). Intriguingly, within the top five most abundantly modified proteins, we noted that ubiquitin was prominently ADP-ribosylated in the presence of PARP10, with over 30-fold induction. Ubiquitin has previously been implicated in the context of PARP10, and is the only PARP enzyme to contain Ubiquitin Interaction Motifs (Verheugd et al., 2013). Thus, we scrutinized the amino acid residues of ubiquitin that were direct targets of ADP-ribosylation, and ten ADPr sites could be quantified on ubiquitin in HeLa cells (Fig. 6D-E). Under standard growth conditions most (∼77%) ADPr was observed near the ubiquitin C-terminus on Arg-74, whereas expression of PARP10 led to ∼86% of the total modification on E-51 (∼57%), D-32 (∼19%), and E-24 (∼10%). From a structural point of view, these three residues are in close proximity and surface-accessible, and located opposite from the di-glycine through which ubiquitin is conjugated (Fig. 6F). Taken together, this indicates that PARP10 is involved in ADP-ribosylation of ubiquitin in the context of immune signaling.

### Interferons drive extensive Glu/Asp/Cys-ADPr on immune-linked networks

A549 cells are commonly used to study immune signaling pathways (Dias et al., 2021; Tang et al., 2023; Wang et al., 2021). We applied our Glu/Asp-ADPr compatible methodology to investigate the ADP-ribosylome in the context of interferon treatment of A549 cells (Fig. 4A). Treatment with either IFNα or IFNγ doubled the number of identified ADPr sites (Fig. 4C and Table S2), while overall abundance increased by 5-fold (Fig. 4D). Volcano plot analysis revealed that upon IFNα and IFNγ treatment, 116 and 146 ADPr target proteins were upregulated, and 5 and 2 were downregulated, respectively (Fig. S7 and Table S3). Of note, both interferon treatments resulted in comparable subsets of proteins being regulated by ADP-ribosylation. Thus, we directly compared proteins modulated in the IFNα and IFNγ experiments (Fig. 7A), which revealed 103 proteins that were significantly regulated by both interferons, with 66 proteins significantly regulated by only one IFN. Out of the 540 ADPr sites identified in A549 cells, 312 (58%) sites resided on the proteins that were regulated in response to both interferons. At the site level, we found that the vast majority of significantly interferon-regulated ADPr sites were either Glu, Asp, or Cys linkages (Fig. 7B). In contrast, Ser-ADPr, even though abundantly present in the samples, was mostly unaffected by interferon treatment.

**Figure 7.**
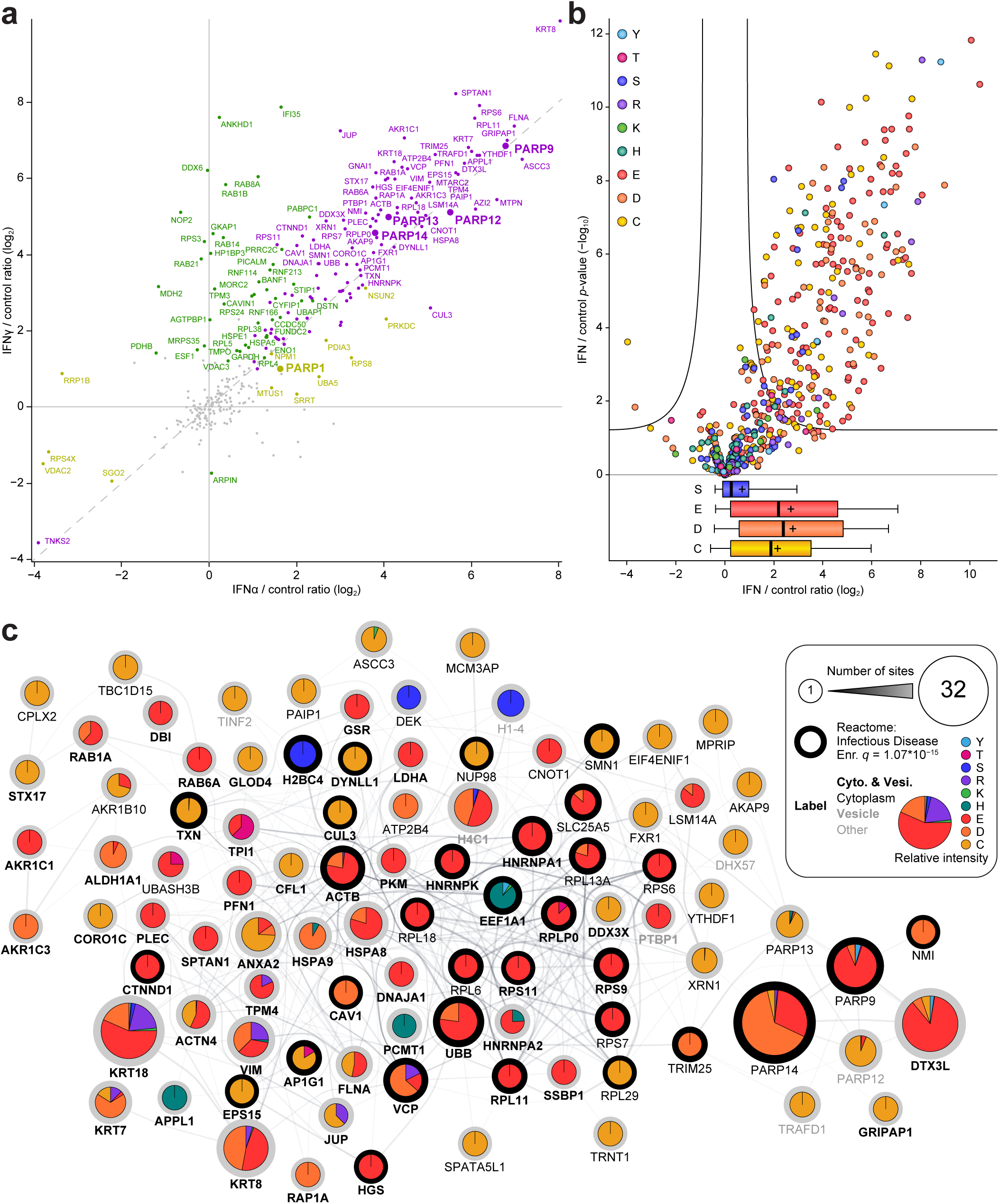
Interferon treatment induces a robust Glu/Asp/Cys ADPr response in A549 cells. (A) Scatter plot analysis, visualizing the ADP-ribosylation ratio of change of target proteins for IFNα (x-axis) and IFNγ (y-axis) treatment of A549 cells, in both cases compared versus untreated cells. Coloring of the proteins and labels indicates statistical significance (*q*<0.01); purple is both, yellow is IFNα, and green is IFNγ. Significance determined via student’s two-sided t-testing with permutation-based FDR control, *n*=4 cell culture replicates. (B) Volcano plot analysis, visualizing ADPr sites stratified by amino acid residue, with the cut-off line at *q*=0.01. Significance determined via student’s two-sided t-testing with permutation-based FDR control, *n*=4 cell culture replicates. The boxplot visualization below summarizes all data points on the x-axis for each amino acid residue. Thick line; median, plus symbol; mean, box limits; 1^st^ and 3^rd^ quartile, whisker limits; 5^th^ and 95^th^ percentile. (C) STRING network analysis of all ADPr target proteins that were significantly upregulated both in response to IFNα and IFNγ. 96 out of 102 proteins were interconnected at default STRING clustering confidence, the 6 disconnected nodes are not shown. Node coloring represents relative contribution of each ADPr amino acid linkage type to overall protein modification, size corresponds to number of sites, label font type indicates cytoplasmic, vesicle, or other localization, and thick node outline indicates the protein is annotated by Reactome as involved in “Infectious Disease”. All annotated terms are significantly enriched within the network, as determined by the STRING database, compared to the total proteome background.

Next, we explored the biological properties of the 102 proteins that were increasingly ADP-ribosylated in A549 cells, both in the case of IFNα and IFNγ treatment (Fig. 7C and Table S3). Via network connectivity analysis using the STRING database (Szklarczyk et al., 2023), 96 out of 102 proteins formed one significantly (*p* < 10^-16^) interconnected cluster, with ADPr on these proteins mainly targeting Glu, Asp, and Cys residues. Interestingly, modification of Cys tended to preclude modification of Glu/Asp, and vice versa. Furthermore, modification by Cys-ADPr seemed to only target one site per protein, whereas Glu/Asp-ADPr frequently targeted up to several dozen sites per protein, in line with observations during our benchmark experiments (Figs. 3A and S3A). Within the interferon-regulated ADPr target network there was an overrepresentation of cytoplasmic proteins, and moreover there was a significant presence of proteins involved in immune regulation, infectious disease, and the cellular response to stress (Table S5). Taken together, our work demonstrates that interferon treatment of A549 cells induces extensive Glu/Asp/Cys-specific ADP-ribosylation of a highly interconnected network of cytoplasmic proteins enriched for immune- and stress-related functions.

### Interferons remodel a core antiviral PARP network through distinct ADPr patterns

We profiled the ADP-ribosylome of A549 cells, as well as HeLa cells expressing PARP10, both treated with various cytokines. Among the hundreds of ADPr target proteins we identified (Table S3), we also found 10 PARP family members to be ADP-ribosylated; PARP1, PARP2, TNKS2, PARP7, PARP8, PARP9, PARP10, PARP12, PARP13, and PARP14 (Fig. 8A). Intriguingly, across two distinct cell lines, and two different interferon treatments, we observed a strong modification of PARP9, PARP12, PARP13, and PARP14. Interestingly, TNKS2 showed an inversed trend in A549 cells following interferon treatment. We also noted PARP10 induction resulting in increased ADP-ribosylation of PARP7 (Fig. 8A), and a stable baseline of PARP1, PARP2, and PARP8 in HeLa cells regardless of treatment.

**Figure 8.**
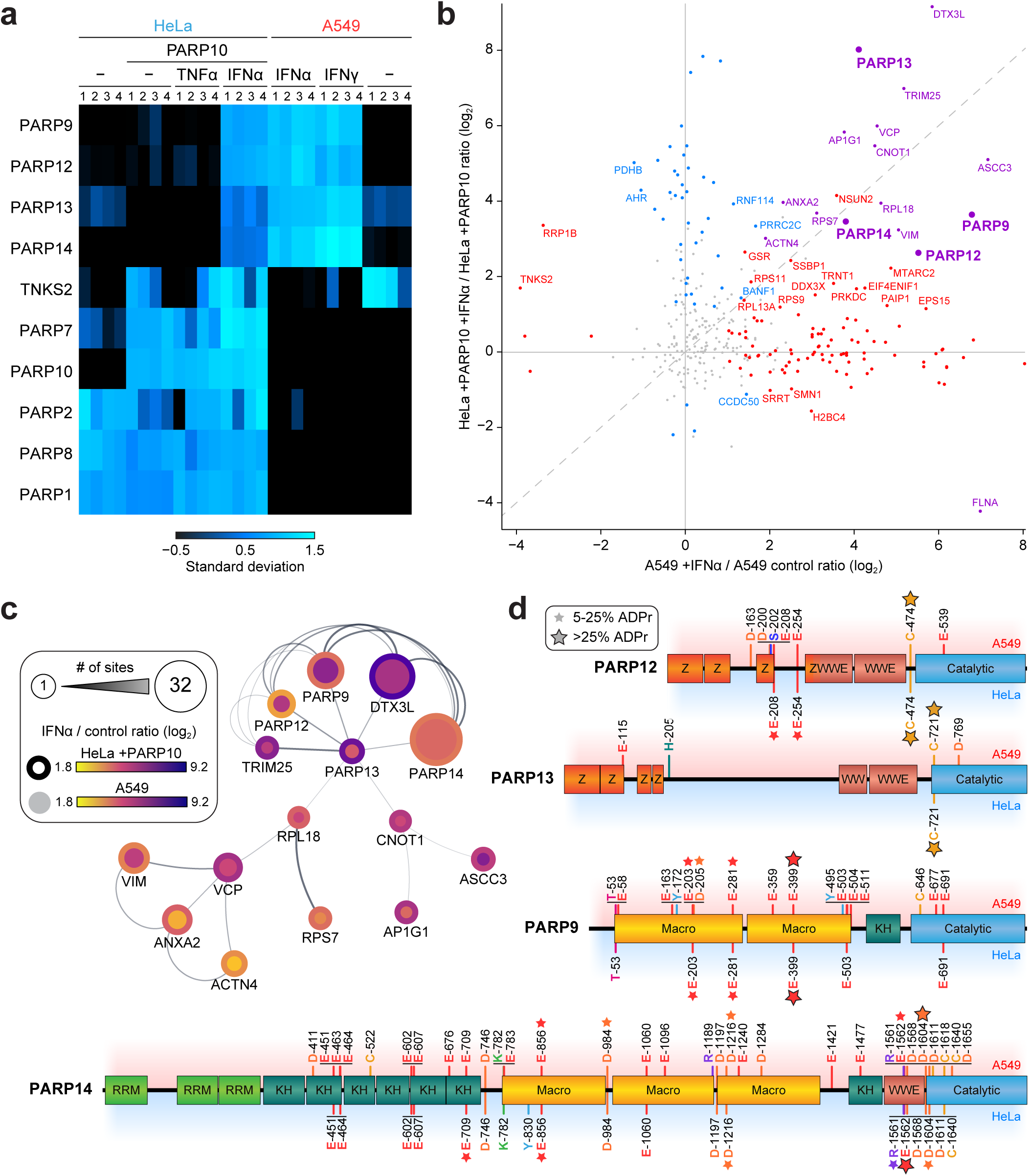
Interferon treatment induces a robust Glu/Asp/Cys ADPr response in A549 cells. (A) Heatmap analysis, displaying how prominently the different PARP family members were detected as ADP-ribosylated across all replicates and conditions. Summed ADPr site abundances were z-scored prior to plotting. (B) Scatter plot analysis, visualizing the ADP-ribosylation ratio of change of ADPr target proteins in response to IFNα treatment in A549 cells (x-axis) and HeLa cells (y-axis), compared versus their respective controls. Coloring of the proteins and labels indicates significance (*q*<0.01); purple is both, red is A549, and blue is HeLa. Significance determined via student’s two-sided t-testing with permutation-based FDR control, *n*=4 cell culture replicates. (C) STRING network analysis of all 15 ADPr target proteins that were significantly upregulated in response to IFNα, both in A549 and HeLa cells. Coloring represents how highly ADPr is induced upon IFNα treatment, with inner node color for A549 and outer node color for HeLa. Size of the nodes scales to the number of ADPr sites per protein. (D) Linear visualization of the PARP proteins from **B** and **C**. Drawings are scaled to the length of the proteins, with known domains indicated. All mapped ADPr sites are marked, stratified by detection in A549 or HeLa. Small stars indicate 5-25% of total protein ADPr detected on the residue, whereas large stars indicate >25% of protein ADPr on the residue.

Comparing the regulation of ADPr target proteins in response to IFNα between A549 cells and HeLa cells expressing PARP10 revealed that the majority of proteins (144; ∼90%) was specifically regulated in either one of the cell lines (Fig. 8B). However, 15 proteins were significantly upregulated in both cell lines upon IFNα treatment (labeled in purple), hinting at a subset of ADPr target proteins that are canonically associated with the cellular immune response (Fig. 8C). Indeed, these proteins formed an interconnected network when clustering via the STRING database, with a sub-cluster including PARP9, PARP12, PARP13, PARP14, DTX3L, and TRIM25, all of which act as key regulators of antiviral and immune signaling pathways (Liu et al., 2025; Luscher et al., 2022b; Malgras et al., 2021). The network also extended into additional cellular processes: transcriptional and RNA regulation (CNOT1, ASCC3), vesicle trafficking (AP1G1), ribosomal translation (RPL18, RPS7), protein quality control (VCP), and cytoskeletal organization and dynamics (VIM, ANXA2, ACTN4).

Having established the interferon-regulated PARP network, we next examined the specific ADPr modification sites within the major antiviral PARPs to resolve how their regulation manifests at the residue level (Fig. 8D). Among the four antiviral PARPs, PARP14 showed the strongest ADP-ribosylation, with a total of 32 sites and predominant Glu/Asp modification. Across both HeLa and A549 cells, consistent ADPr modification was observed at E-856 and D-1216 in the PARP14 macrodomains, and at E-1562 and D-1604 in the WWE domain near the interface with the catalytic domain. The second-most modified antiviral PARP family member, PARP9, was modified on 16 sites. The majority of ADPr occurred on E-203, E-281, E-399, all residing in the macrodomains. The remaining two antiviral PARPs; PARP12 and PARP13, were not as heavily ADP-ribosylated, and interestingly both PARPs were predominantly modified on a single Cys residue, C-474 and C-721 respectively, located at the N-terminus of their catalytic domains.

In contrast to the antiviral PARPs, the other PARP family members displayed different site preferences. As anticipated, despite PARP1 modification occurring on a multitude of Glu/Asp residues, the vast majority of intensity (∼99% in HeLa and ∼96% in A549) was observed on Ser residues (Fig. S8). Reassuringly, the Glu/Asp ADPr sites we detected here in PARP1 overlap well with the Glu/Asp sites reported by the Matic lab (Longarini and Matic, 2024), and in both cases the ADPr resided in the auto-modification domain and towards the C-terminus of the second zinc finger domain. Expectedly, PARP2 ADPr exclusively occurred on a Ser residue. For the remaining four PARP family members, TNKS2, PARP7, PARP8, and PARP10, we were only able to detect between one and three ADPr sites, and interestingly in all cases >75% of modification occurred on Cys residues.

Taken together, our findings highlight a distinct subset of antiviral PARPs and associated proteins that are consistently ADP-ribosylated across cell types in response to interferon treatment, forming a core immune-regulated network. Moreover, our site-specific profiling of ADPr sites reveals distinct modification patterns across PARP family members and other immune system proteins, and represents a valuable resource for the field.

## DISCUSSION

In this work, we augmented our established Af1521 proteomics enrichment workflow to profile glutamate- and aspartate-linked ADP-ribosylation, and applied it to cytokine-stimulated A549 and HeLa cells expressing PARP10. We profiled >600 Glu/Asp-ADPr sites across diverse cellular contexts, revealing extensive cell type- and treatment-specific ADPr patterns, but also uncovering a conserved interferon-induced network centered on antiviral PARPs and immune regulators.

The lability of ester-linked Glu/Asp-ADPr has recently been established (Javed et al., 2023; Longarini and Matic, 2024; Tashiro et al., 2023), with the main drivers of lability being alkaline sample preparation conditions along with elevated temperatures and extended incubation times. Previously, our lab performed a range of MS-based proteomics studies wherein the Af1521 methodology was applied using alkaline conditions (Hendriks et al., 2021; Larsen et al., 2018), and similarly other labs also predominantly use similar conditions that are not necessarily compatible with the total preservation of Glu/Asp-ADPr (Bilan et al., 2017; Zhang et al., 2013). Thus, we and others may have underestimated the prevalence of Glu/Asp-ADPr in the past, which should be taken into account when assessing previous datasets. However, the context of the experiments remains important, as the majority of our studies revolved around induction of DNA damage via treatment with oxidizing or alkylating agents (Buch-Larsen et al., 2021; Hendriks et al., 2019; Larsen et al., 2018), which primarily trigger a robust Ser-ADPr response which is accurately visualized using Af1521 under both alkaline and acidic conditions (Fig. 2). Importantly, it should also be mentioned that a relevant biological system and context is required for studying Glu/Asp-ADPr, as this linkage type usually requires activation of a specific subset of writer enzymes (Luscher et al., 2018), or otherwise has mainly been observed in HPF1-deficient cell systems (Bonfiglio et al., 2017; Leung, 2017; Longarini and Matic, 2024).

Another relevant aspect to consider is the hydrolase activity of Af1521 itself (Jankevicius et al., 2013; Rosenthal et al., 2013). Although there is some irony in using an active eraser enzyme to purify its target modification, we have demonstrated in the past that Af1521 is exceedingly efficient in enriching the majority of ADPr linkage types (Hendriks et al., 2019), and within the systems investigated we found no evidence of Af1521 reversing ADPr during the purification (Jungmichel et al., 2013). However, the hydrolase activity of Af1521 targeting Glu/Asp-ADPr, along with alkaline conditions employed during the ADPr enrichment procedure, could surely result in abolishment of Glu/Asp-linked ADPr during sample preparation. In this work, we circumvent the reversal of Glu/Asp-ADPr by using acidic conditions, lower temperatures, and shorter incubation times. Moreover, we found that the mutant Af1521, which has been shown to most efficiently enrich Ser-ADPr (Nowak et al., 2020), is either more efficient at stably binding Glu/Asp-ADPr, or is diminished in its ability to hydrolyze Glu/Asp-ADPr.

We observed a notable number of Cys-ADPr sites across our experiments. Specifically, in the benchmarking experiment (Fig. 2), Cys-ADPr was observed predominantly when using the mutant Af1521 to enrich ADPr, and with a preference for alkaline conditions. Since Cys-ADPr has been reported to occur enzymatically and non-enzymatically (McDonald and Moss, 1994), it could be speculated that under alkaline conditions Glu/Asp-ADPr may non-enzymatically transfer to Cys residues. However, when comparing A549 cells treated with interferon and U2OS HPF1 knockout cells treated with H_2_O_2_, we found a notably different ratio between Glu/Asp and Cys-ADPr observed under acidic and alkaline conditions, whereas a similar ratio would be expected in case of non-enzymatic transfer. Further, we investigated the sequence context of ADP-ribosylated and unmodified Cys residues, and found no significant difference (*p* = 0.36) between proximal (±7 aa) occurrences of Glu/Asp residues, with on average 1.75 Glu/Asp residues nearby 209 Cys-ADPr sites, and on average 1.83 Glu/Asp residues nearby 2,540 unmodified Cys residues from the same proteins (Table S6). Thus, based on these two distinct observations, the Cys-ADPr we mapped likely has a biological origin. Furthering this, we observed Cys-ADPr to be the predominant (>2/3^rd^) modification on 119 proteins (Table S3), including PARP7/8/10/12/13, TNKS2 and TNKS1BP1, as well as RNF166 which has been reported to control TNKS stability (Perrard and Smith, 2023). Additionally, 43 of the Cys-ADPr target proteins are annotated to have RNA binding properties (Table S3), which corroborates a previous proteome-wide analysis that demonstrated PARP7 catalyzing Cys-ADPr, with many of its targets involved in RNA binding and other RNA regulatory roles (Rodriguez et al., 2021). Collectively, these findings support Cys-ADPr as a biologically relevant modification with potential roles in RNA regulation and protein stability.

Glu/Asp-ADPr has historically been profiled using hydroxylamine derivatization of the ADPr moiety, typically following boronate affinity enrichment (Gibson et al., 2016; Zhang et al., 2013; Zhen et al., 2017). Recently, the Matic lab published an antibody-based strategy for ADPr enrichment, wherein they also employed acidic conditions to stabilize Glu/Asp-ADPr (Longarini and Matic, 2024). Based on our comparison of ADPr sites and target proteins across all studies, we found that we best overlapped with the larger hydroxylamine studies at the target protein level, whereas at the site-specific level the highest agreement was with the recent study from the Matic lab (Fig. 5C). We reason that the protein-level overlap could align with the overall number of ADPr target proteins identified, with the overlaps correlating with the size of the studies. At the site-specific level, both technical and biological factors are likely to play a role. First, we detect the full ADPr moiety using ETD-based fragmentation similar to the Matic lab, whereas other studies detect a hydroxylamine derivative using HCD-based fragmentation. Second, whereas the hydroxylamine-based method exclusively detects Glu/Asp-ADPr, the approach presented in this work is capable of detecting any ADPr linkage type. Third, in this study we employed acidic conditions during sample preparation, whereas in the hydroxylamine studies the samples are often first processed under alkaline conditions, which could result in some loss of Glu/Asp-ADPr prior to its reaction with hydroxylamine. Fourth, the Matic lab investigated Glu/Asp-ADPr in the absence of HPF1 to increase the cellular levels of Glu/Asp-ADPr (Palazzo et al., 2018), which aligns with our benchmark experiment, and generated the largest overlap, likely due to similarity of the biological context studied. Overall, considering the promiscuity of ADPr and its plethora of biological functions, and further convolution by differences in technical methodology, a large degree of heterogeneity is to be expected across different studies.

Our results reveal that PARP10 expression induces a profound rewiring of the ADP-ribosylome, shifting modification patterns from nuclear toward cytosolic proteins, in line with the cytoplasmic localization and signaling roles of PARP10 described previously (Kleine et al., 2012). Prior to PARP10 expression, we identify ubiquitin as a prominent ADPr target protein, with modification mainly occurring on Arg-74 adjacent to the canonical di-glycine conjugation site. PARP9/DTX3L was previously found to drive ADPr to the C-terminus of ubiquitin (Yang et al., 2017). Our mass spectrometry mainly support ADPr on Arg-74, with sparse to no evidence supporting modification of the C-terminus. Nevertheless, we find DTX3L to become less ADP-ribosylated upon PARP10 expression, along with a shift from modification near the C-terminus of ubiquitin to multiple acidic residues that cluster in a structurally accessible region opposite the di-glycine, suggesting that DTX3L modulates ubiquitin ADPr under baseline conditions.

IFNα and IFNγ regulate distinct and overlapping sets of proteins (Megger et al., 2017), and we found that in A549 cells the ADP-ribosylome was similarly modulated by both interferons, in either case triggering an extensive remodelling of the ADP-ribosylome, with a roughly twofold increase in the number of identified sites and a fivefold increase in overall ADPr abundance. At the site-specific level, we observed that the interferon-regulated ADPr was dominated by Glu, Asp, and Cys linkages, whereas Ser-ADPr remained largely unaffected, suggesting a preferential engagement of Glu/Asp/Cys-ADPr in interferon-responsive pathways. Network analysis revealed that the majority of interferon-regulated ADPr targets formed a highly interconnected cluster of cytoplasmic proteins enriched for immune, antiviral, and stress-related functions. Within this network, we identified a prominent axis of antiviral PARPs and associated factors, including PARP9, PARP14, and DTX3L, with additional contributions from PARP12, PARP13, and TRIM25. Interferon treatment is known to activate PARP14, which then becomes the main driver of mono-ADPr in response, with PARP9/DTX3L regulating the activity of PARP14 (Kar et al., 2024; Ribeiro et al., 2024). Consistently, in our data, these three proteins are the most abundantly ADP-ribosylated, with dozens of Glu/Asp modification sites. Immunoprecipitation of p62 from IFNγ-treated cells followed by mass spectrometry revealed interaction with and regulation of PARP14, PARP9, and DTX3L (Raja et al., 2025), along with several dozen other proteins for which we mapped ADPr sites in our screen. Although it is not clear from our data which PARPs target the aforementioned proteins, it is known that PARP14 targets p62, PARP13, RNF114, and RNF166 (Dukic et al., 2023). Altogether, the coordinated ADPr of these proteins highlights a conserved, interferon-responsive module that may act as a central hub for antiviral signaling, linking residue-specific ADPr modifications to functional regulation of the innate immune response.

In conclusion, our work highlights the relevance of tailoring mass spectrometry-based workflows to the specific chemistry of ADP-ribosylation. As demonstrated here, subtle differences in buffer conditions, temperature, incubation times, and enrichment reagents during sample preparation can strongly influence the detection and quantification of labile ADP-ribosylation linkages, particularly on Glu/Asp residues. No single method is universally optimal, and differences in sample preparation and fragmentation strategies likely contribute to the heterogeneity observed across studies. Using an optimized acidic workflow with a mutant Af1521 macrodomain and ETD-based fragmentation, we achieved deep, site-specific coverage of Glu/Asp-ADPr across multiple cellular contexts and cytokine treatments. This dataset provides a rich resource for functional studies or mutagenesis, highlights both the context-specific nature of ADP-ribosylation and conserved networks such as the interferon-responsive antiviral PARP axis, while serving as a methodological reference for future ADPr profiling.

## METHODS

### Cell lines and cell culture

HeLa cells (CCL-2, female) and A549 cells (CCL-185, male) were acquired via the American Type Culture Collection. U2OS cells with HPF1 knockout (ΔHPF1) (Gibbs-Seymour et al., 2016) and HeLa cells with inducible PARP10 expression (Herzog et al., 2013), were described previously. All cells were cultured at 37 °C and 5% CO2 in Dulbecco’s Modified Eagle’s Medium (Invitrogen) supplemented with 10% fetal bovine serum and a penicillin/streptomycin mixture (100 U/mL; Gibco). All cells were routinely tested for mycoplasma. Cells were not routinely authenticated.

### Cell treatment

For induction of ADP-ribosylation in HeLa cells and U2OS ΔHPF1 cells, treatment was performed with 1 mM H_2_O_2_ (Sigma Aldrich) for 10 min or 30 min, respectively. For induction of ADPr signaling in A549 cells, treatment was performed for 24 h with either 0.684 ng/mL (∼180 U/mL) IFNα (Peprotech) or 100 ng/mL (∼194 U/mL) IFNγ (Peprotech). To express PARP10 in inducible HeLa cells (Herzog et al., 2013), 1 µg/mL doxycycline was added for 24 h. Concomitantly, ADPr signaling was induced in the cells via addition of 0.684 ng/mL (∼180 U/mL) IFNα (Peprotech) for 24 h, or addition of 5 ng/mL (∼196 U/mL) TNFα (Peprotech) during the last 6 h. Approximately 200 million cells (∼3 confluent square 25-cm dishes) were cultured per replicate. For downstream processing and MS analysis, replicate batches were divided over multiple purification conditions and technical replicates.

### Cell lysis and protein digestion

For analysis of ADPr at alkaline pH, the full procedure for proteolytic digestion of cells was done mostly as described previously (Hendriks et al., 2019; Larsen et al., 2018). For purification of ADPr at acidic pH, the protocol was adapted considerably, with all changes outlined here. Cells were washed once with ice-cold PBS, and gently scraped at 4 °C in a minimal volume of PBS. For benchmarking experiments, all cell batches were divided equally between two tubes prior to lysis in different buffers. For quantitative experiments, all replicates were subjected to the acidic workflow. Cells were pelleted by centrifugation at 500*g*, and either lysed in 10 pellet volumes of Alkaline Lysis Buffer (6 M guanidine-HCl, 50 mM Tris-HCl, pH 8.5), or 10 pellet volumes of Acidic Lysis Buffer (6 M guanidine-HCl, 10 mM Tris-HCl, 100 mM sodium phosphate, pH 6.3). Complete lysis was achieved by alternating vigorous shaking with vigorous vortexing, for 30 seconds, prior to snap freezing of the lysates using liquid nitrogen. Frozen lysates were stored at −80 °C until further processing. Lysates were thawed and sonicated at 30 W, for 1 second per 1 mL of lysate, spread across 2 separate pulses. For acidic samples, care was taken to keep the lysates below 30 °C. Tris(2-carboxyethyl)phosphine and chloroacetamide were added to final concentrations of 10 mM. For alkaline samples, proteins were digested using Lysyl Endopeptidase (Lys-C, 1:200 w/w; Wako Chemicals) for 3 hours at 25 °C. For acidic samples in the benchmark experiment, proteins were digested using Lys-C (1:100 w/w) for 1 hour at 25 °C. For acidic samples in the quantitative experiment, lysates were first diluted to a final concentration of 3 M guanidine-HCl by addition of 100 mM sodium phosphate pH 6.3, prior to digestion using Lys-C (1:150 w/w) for 1 hour at 25 °C. Following Lys-C digestion, alkaline samples were diluted to a final concentration of 1.5 M guanidine-HCl using 50 mM Tris-HCl pH 8.5, and acidic samples were diluted to a final concentration of 1.5 M guanidine-HCl using 100 mM sodium phosphate pH 6.3. Alkaline samples were further digested overnight using modified sequencing grade Trypsin (1:200 w/w; Sigma Aldrich). Acidic samples were further digested using sequencing grade Trypsin (1:150 w/w) for 1 hour at 25 °C. Following digestion, samples were (further) acidified by addition of trifluoroacetic acid (TFA) to a final concentration of 0.5% (v/v), cleared by centrifugation, and purified using reversed-phase C18 cartridges (Sep-Pak C18 6 cc Vac Cartridge, 500 mg Sorbent, Waters) according to the manufacturer’s instructions, and using one cartridge per 20 mg of input protein. Elution of peptides was performed with 35% acetonitrile (ACN) in 0.1% TFA, peptides were frozen overnight at −80 °C, and afterwards lyophilized for 96 h.

### Purification of ADP-ribosylated peptides using Af1521

Lyophilized peptides were dissolved in either Alkaline AP buffer (50 mM Tris-HCl, 1 mM MgCl_2_, 250 μM DTT, and 50 mM NaCl, pH 8.0) or Acidic AP buffer (10 mM Tris-HCl, 100 mM sodium phosphate, 1 mM MgCl_2_, 250 μM DTT, and 50 mM NaCl, pH 6.3). Dissolved peptides were cleared by centrifugation and supernatant decanted into clean tubes. Subsequently, peptides were either left untreated or treated with PARG enzyme (kind gift from Prof. Dr. Michael O. Hottiger), with PARG added in a 1:1,000 (w/w) ratio for 1 h at room temperature. GST- or His10-tagged Af1521 macrodomain was produced in-house using BL21(DE3) bacteria, and coupled to glutathione Sepharose 4B beads (Sigma-Aldrich) or Ni-NTA Agarose beads (QIAGEN), respectively. Bacterial production of Af1521 was performed essentially as described previously (Hendriks et al., 2019; Larsen et al., 2018), although for His10-tagged Af1521 an EDTA-free Protease Inhibitor Cocktail (Sigma-Aldrich) was used. Both wildtype Af1521 and Af1521-K35E/Y145R (Nowak et al., 2020) were produced with both epitope tags. For benchmarking experiments, Sepharose beads with GST-tagged Af1521 or Af1521-K35E/Y145R were used for alkaline samples, whereas Agarose beads with His10-tagged Af1521 or Af1521-K35E/Y145R were used for acidic samples. For quantitative experiments, the (acidic) samples were split equally and purified using either His10- or GST-tagged Af1521-K35E/Y145R. In all cases, peptides were chilled to 4 °C prior to addition of beads with Af1521, and otherwise kept on ice or inside a cold room (4 °C) at all points during the purification. The bead mixture was allowed to incubate on a rotating mixer at 4 °C for 45 min. For alkaline samples, beads were sequentially washed two times with ice-cold Alkaline AP Buffer, two times with ice-cold PBS, and two times with ice-cold MQ water. For acidic samples, beads were sequentially washed four times with ice-cold Acidic AP Buffer. On the first wash, beads were transferred to 1.5 mL protein-LoBind tubes (Eppendorf), and LoBind tubes were exclusively used from this point on to minimize loss of peptide. Following the third wash, beads were transferred to a fresh LoBind tube. For samples during the benchmarking experiments, ADP-ribosylated peptides were eluted from the beads using two elution steps of 15 min each, with two bead volumes ice-cold 0.15% TFA, and the pooled elutions were cleared through 0.45 μm spin filters (Ultrafree-MC, Millipore) and subsequently through pre-washed 100 kDa cut-off filters (Vivacon 500, Sartorius). For acidic samples during the benchmarking experiments, ADP-ribosylated peptides were additionally passed through pre-washed 10 kDa cut-off filters (Vivacon 500, Sartorius). For acidic samples during the quantitative experiments, half of the samples were purified using His10-Af1521-K35E/Y145R, with the other half purified using GST-Af1521-K35E/Y145R. Elution from the His10 Agarose beads was performed using two steps of 15 min each, using two bead volumes ice-cold 0.15% TFA, with the pooled elutions passed through 0.45 μm, 100 kDa, and 10 kDa filters as described above. The cleared elutions were split in two halves, of which one set of halves were left as is, whereas the other set of halves were pH-neutralized via addition of 10 volumes of ice-cold 200 mM sodium phosphate pH 6.3, and cleared from histidine-rich background binders via a 15 min incubation with empty Ni-NTA beads, prior to re-acidification by addition of TFA to a final concentration of 1%. Elution from GST Agarose beads was performed firstly using two steps of 15 min each, with two bead volumes of ice-cold 0.1% formic acid (FA), and secondly for 15 min using four bead volumes of ice-cold 0.15% TFA. The FA and TFA elutions were kept separately, and both passed through 0.45 μm and 100 kDa filters as described above. After filtering, the FA elution was further acidified by addition of TFA to a final concentration of 0.15%.

### StageTip purification of peptides

The filtered ADP-ribosylated peptides were purified using C18 StageTips at low pH. To this end, C18 StageTips were prepared in-house, by layering four plugs of C18 material (Sigma-Aldrich, Empore SPE Disks, C18, 47 mm) per StageTip. Activation of StageTips was performed with 100 μL 100% methanol, followed by equilibration using 100 μL 80% ACN in 0.1% FA, and two washes with 100 μL 0.1% FA. Samples that were not already acidic (pH < 3) were acidified via addition of TFA to a final concentration of 1%. Samples were centrifuged for 5 min at 14,000g, after which supernatants were loaded on StageTips. Subsequently, StageTips were washed twice using 100 μL 0.1% FA, after which peptides were eluted using 80 µL 30% ACN in 0.1% formic acid. All fractions were dried to completion using a SpeedVac at 60 °C. Dried peptides were dissolved in 20 μL 0.1% FA and stored at −20 °C until MS analysis.

### Peptide synthesis

Peptides containing ADPr on specific moieties were generated essentially as described previously (Tashiro et al., 2022).

The following peptides were synthesized:

NH2-GWTARKSAEAGTAGK-amidated, either unmodified or with ADPr on Arg-5

NH2-GWTARKSAAAGTAGK-amidated, either unmodified or with ADPr on Ser-7

NH2-GWTARKAAEAGTAGK-amidated, either unmodified or with ADPr on Glu-9

NH2-GWTARKAADAGTAGK-amidated, either unmodified or with ADPr on Asp-9

Peptide sequences were chosen to have favorable properties for electron-transfer dissociation-based fragmentation, via inclusion of several arginine and lysine residues to ensure a suitable charge state during positive mode mass spectrometric analysis. Peptide sequences were otherwise kept as similar as possible to reduce bias, but nonetheless were not fully isobaric so that peptides may easily be distinguished from a mixture.

### Synthetic peptide treatment

All lyophilized peptides were dissolved in 0.1% FA, and investigated using mass spectrometry for their overall ionization efficiencies (i.e. perceived signal by the mass spectrometer). A mixture with comparable amounts of all four ADP-ribosylated peptides was generated based on these observations, and used for all experiments. The synthetic ADPr peptide mixture was treated in various ways, as outlined in the figure legends. The “control” condition for all experiments corresponds to an otherwise untreated ADPr peptide mixture. Treatment at different pH utilized the following buffers. For “pH 3”, 0.1% formic acid. For “pH 5”, sodium acetate and acidic acid mixture calibrated to pH 5.0. For “pH 6”, 100 mM sodium phosphate calibrated to pH 6.3. For “pH 7”, phosphate buffered saline pH 7.2. For “pH 8.5”, Tris-HCl calibrated to pH 8.5. For “pH 11”, 50 mM ammonium hydroxide. For all non-control conditions from the experiments described in Figure 1C, 1D, S1A, and S1B, and following their treatments, all peptide mixtures were acidified using 1% TFA (v/v) and purified using C18 StageTip as described above.

### MS dataset overview

Dataset 1: Synthetic peptide experiments, related to Figure 1, referred to as “DS1”.

Dataset 2: Benchmarking experiments, related to Figures 2 and 3, referred to as “DS2”.

Dataset 3: Quantitative experiments, related to Figures 4 through 8, referred to as “DS3”.

### ETD-based mass spectrometric analysis

All samples were analyzed using an Orbitrap Fusion™ Lumos™ Tribrid™ mass spectrometer (Thermo), and analyzed on 15-cm long analytical columns with an internal diameter of 75 μm, packed in-house using ReproSil-Pur 120 C18-AQ 1.9 µm beads (Dr. Maisch). On-line reversed-phase liquid chromatography to separate peptides was performed using an EASY-nLC™ 1200 system (Thermo), and the analytical column was heated to 40°C using a column oven (Sonation). For measurement of DS3 samples, n-dodecyl β-D-maltoside (DDM) was added to a final concentration of 0.005%. Peptides were eluted from the column using a gradient of Buffer A (0.1% formic acid) and Buffer B (80% ACN in 0.1% FA). For DS1, the primary gradient ranged from 3% B to 38% B over 18 minutes, followed by a washing block of 7 minutes. For DS2 and His10-Af1521 samples from DS3, the primary gradient ranged from 3% B to 24% B over 50 min, followed by a washing block of 30 min, with the times halved for a second acquisition replicate. For GST-Af1521 samples from DS3, the primary gradient ranged from 4% B to 40% B over 35 min, followed by a washing block of 15 min. Electrospray ionization (ESI) was achieved using a Nanospray Flex Ion Source (Thermo). Spray voltage was set to 2 kV, capillary temperature to 275°C, and RF level to 40%. Full scans were performed at a resolution of 120,000, with a scan range of 300 to 1,300 m/z (DS1,2) or 350 to 1,250 (DS3), a maximum injection time of 50 ms, and an automatic gain control (AGC) target of 600,000 (DS1,2) or 1,000,000 (DS3) charges. Precursors were isolated from the entire full scan range (DS1) or a range of 325 to 1,275 (DS2) or 400 to 1,200 m/z (DS3). Isolation was performed at a width of 1.3 m/z, an AGC target of 200,000 charges, and precursor fragmentation was accomplished using electron transfer disassociation with supplemental higher-collisional disassociation (EThcD) at 20 NCE, using calibrated charge-dependent ETD parameters (Rose et al., 2015). Precursors with charge state 2-6 (DS1) or 3-5 (DS2,3) were isolated for MS/MS analysis, and for DS2 and DS3 prioritized from charge 3 (highest) to charge 5 (lowest). Selected precursors were excluded from repeated sequencing by setting a dynamic exclusion of 0.6 s per minute (i.e. 1%) of total gradient time. MS/MS spectra were measured in the Orbitrap at a scan resolution of 60,000, and at various levels of sensitivity, by using TopN and maximum MS2 injection time combinations of: Top5 with 120 ms, Top4 with 180 ms, Top3 with 250 ms, and Top2 with 500 ms.

### Synthetic peptide mass spectrometry data analysis

For DS1, analysis of the mass spectrometry raw data was performed using Skyline software (Pino et al., 2020). To this end, Skyline was provided with monoisotopic peptide information corresponding to each of the eight peptides, obtained via a pre-search of representative data files using MaxQuant software (Cox and Mann, 2008; Cox et al., 2011), version 1.5.3.30. C-terminal amidation was set as a fixed peptide modification. ADP-ribosylation was set as a variable modification, as well as tryptophan oxidation to kynurenin, as we observed this to be highly prevalent. Thus, Skyline was set up to identify and extract intensity profiles for the following 16 peptide sequences directly from the data.

GWTARKSAEAGTAGK[−1], GW[+4]TARKSAEAGTAGK[−1], GWTAR[+541.1]KSAEAGTAGK[−1],

GW[+4]TAR[+541.1]KSAEAGTAGK[−1], GWTARKSAAAGTAGK[−1],

GW[+4]TARKSAAAGTAGK[−1], GWTARKS[+541.1]AAAGTAGK[−1],

GW[+4]TARKS[+541.1]AAAGTAGK[−1], GWTARKAAEAGTAGK[−1],

GW[+4]TARKAAEAGTAGK[−1], GWTARKAAE[+541.1]AGTAGK[−1],

GW[+4]TARKAAE[+541.1]AGTAGK[−1], GWTARKAADAGTAGK[−1],

GW[+4]TARKAADAGTAGK[−1], GWTARKAAD[+541.1]AGTAGK[−1], and

GW[+4]TARKAAD[+541.1]AGTAGK[−1].

Extracted peptide intensities were normalized to total ion current, and variants of tryptophan oxidation were summed. To calculate the fraction of ADP-ribosylated peptide for each amino acid linkage, the intensity of their respective modified peptide was divided by the summed intensity of both unmodified and modified peptides.

### ADP-ribosylome mass spectrometry data analysis

For DS2 and DS3, analysis of the mass spectrometry raw data was performed using MaxQuant software (Cox and Mann, 2008; Cox et al., 2011), version 1.5.3.30. MaxQuant default settings were used, with exceptions outlined below. Two separate computational searches were performed, one for each separate MS dataset (DS2,3). For generation of the theoretical spectral library, a HUMAN.fasta database was downloaded from UniProt on the 29^th^ of April, 2023. For DS3, the sequence for the Af1521 macrodomain was also included in the search. N-terminal acetylation, methionine oxidation, cysteine carbamidomethylation, and ADP-ribosylation on all amino acid residues known to potentially be modified (C, D, E, H, K, R, S, T, and Y), were included as variable modifications. For the first search, which is only used for mass recalibration, no variable modifications were used. Up to 8 missed cleavages were allowed, a maximum allowance of 3 variable modifications per peptide was used, and maximum peptide mass was set to 6,000 Da. Second peptide search was enabled for DS2 only. Matching between runs was enabled with an alignment time window of 20 min, and a match time window of 60 s. First and main search precursor mass tolerances of 10 and 4.5 ppm were used, respectively. For fragment ion masses, a tolerance of 20 ppm was used. Modified peptides were filtered to have an Andromeda score of >40 (default), and a delta score of >20. A site decoy fraction of 2% was applied. Data was automatically filtered by posterior error probability to achieve a false discovery rate of <1% (default), at the peptide-spectrum match and the protein assignment levels.

### Non-specific data search

For DS2, a non-specific data search was performed wherein ADPr is allowed to reside on any of the 20 naturally occurring amino acids, for purposes of validating amino acid specificity and localization potential within the MS data. MaxQuant was used essentially as described above, with the following changes in settings. Maximum missed cleavages were reduced to 4. Only ADP-ribosylation on all 20 amino acids was set as a variable modification, with a maximum of one variable modification per peptide. Maximum peptide mass was restricted to 2,500 Da. Second peptide search was disabled. The “evidence.txt” output file from MaxQuant was used to extract raw localization probabilities.

### Data filtering

Beyond automatic filtering and FDR control as applied by MaxQuant, the data were manually stringently filtered in order to ensure proper identification and localization of ADP-ribose. PSMs modified by more than one ADP-ribose were omitted. PSMs corresponding to unique peptides were used for ADP-ribosylation site assignment if localization probability was >0.75. Erroneous MaxQuant intensity assignments were manually corrected in the sites table, and based on localized PSMs only. For ADP-ribosylation target proteins, a list was generated solely based on ADP-ribosylation sites, with multiple sites within the same protein summed up for quantification purposes.

### Comparison to other ADPr MS studies

In order to align ADPr target proteins, a scaffold was created based on all unique protein-coding genes as downloaded from UniProt on the 29^th^ of April, 2023. ADPr target proteins identified in this study were aligned to the scaffold, along with ADPr target proteins identified in four other ADPr proteomics studies (Gibson et al., 2016; Longarini and Matic, 2024; Zhang et al., 2013; Zhen et al., 2017). Alignment was primarily performed using Uniprot identifier, and otherwise protein-coding gene. For comparison of ADPr sites, a scaffold was generated from all 51 amino acid sequence windows derived from this study, as well as from 51 amino acid sequence windows generated from sites identified in four other ADPr proteomics studies (Gibson et al., 2016; Longarini and Matic, 2024; Zhang et al., 2013; Zhen et al., 2017). For the three older studies, since the reported site information did not align with the current version of UniProt, the reported sequence information was re-aligned to the up-to-date UniProt human FASTA file prior to extraction of 51 amino acid sequence windows. Any duplicate 51 amino acid sequences were excluded, and afterwards all studies were individually mapped to the scaffold.

### Quantification and statistical analysis

Details regarding the statistical analysis can be found in the respective figure legends. All quantitative experiments were performed using at least four replicates to ensure sufficient statistical power. Statistical handling of the data was primarily performed using the freely available Perseus software (Tyanova et al., 2016), and includes term enrichment analysis through FDR-controlled Fisher Exact testing, scatter plot analysis and linear regression, volcano plot analysis (using permutated-based FDR control and an s0 value of 0.5), hierarchical clustering, and principle component analysis. Visualization of the Af1521 structure was performed using ChimeraX (Meng et al., 2023). Venn diagram overlaps were performed using Venny (https://bioinfogp.cnb.csic.es/tools/venny/), and scaled Venn diagrams were rendered using eulerAPE (https://www.eulerdiagrams.com/eulerAPE/v2/). UpSet plots were generated using ChiPlot (https://www.chiplot.online/). Boxplots were drawn using BoxPlotR (http://shiny.chemgrid.org/boxplotr/). Interconnected protein networks were acquired using the STRING database (Szklarczyk et al., 2019), and the layout was designed using Cytoscape (Shannon et al., 2003). Plotting of graphs was done using either Perseus, GraphPad Prism, or Microsoft Excel. All art was drawn and all graphs were polished using Adobe Illustrator.

## DATA AVAILABILITY

The mass spectrometry proteomics data generated in this study have been deposited in the ProteomeXchange Consortium via the PRIDE (Perez-Riverol et al., 2019) partner repository, under accession codes:

## INSERT DATA HERE

All other data generated in this study are provided in the Supplementary Information/Source Data file. Source data are provided with this paper.

## ACKNOWLEDGEMENTS

The work carried out in this study was in part supported by the Novo Nordisk Foundation Center for Protein Research, the Novo Nordisk Foundation (grant agreement numbers NNF14CC0001, NNF13OC0006477, and NNF24SA0098829), Danish Council of Independent Research (grant agreement numbers 8020-00220B and 0135-00096B), The Danish Cancer Society (grant agreement R146-A9159-16-S2), and by a center-of-excellence grant from the Danish National Research Foundation to Copenhagen Center for Glycocalyx Research (DNRF196). We would like to thank members of the NNF-CPR Mass Spectrometry Platform for instrument support and technical assistance, and the lab of Michael O. Hottiger (University of Zurich) for the expression and purification of recombinant human PARG.

## AUTHOR CONTRIBUTIONS

S.C.B-L., I.A.H, I.A, and M.L.N. conceived the project. K.T. and G.L. prepared synthetic ADPr peptides. S.C.B-L. and I.A.H. prepared all samples, measured all samples on the mass spectrometer (MS), processed all MS raw data, and performed MS data analysis. J.D.E. assisted with MS data analysis. S.C.B-L. and I.A.H. prepared all figures and wrote the first draft of the manuscript. B.L. supplied PARP10-inducible cells and provided consult. S.Y.V., J.V.O., and M.L.N. provided mass spectrometer infrastructure. S.C.B-L., I.A.H., and I.A. finalized the manuscript with input from all authors.

## SUPPLEMENTARY FIGURE LEGENDS

**Figure S1.**
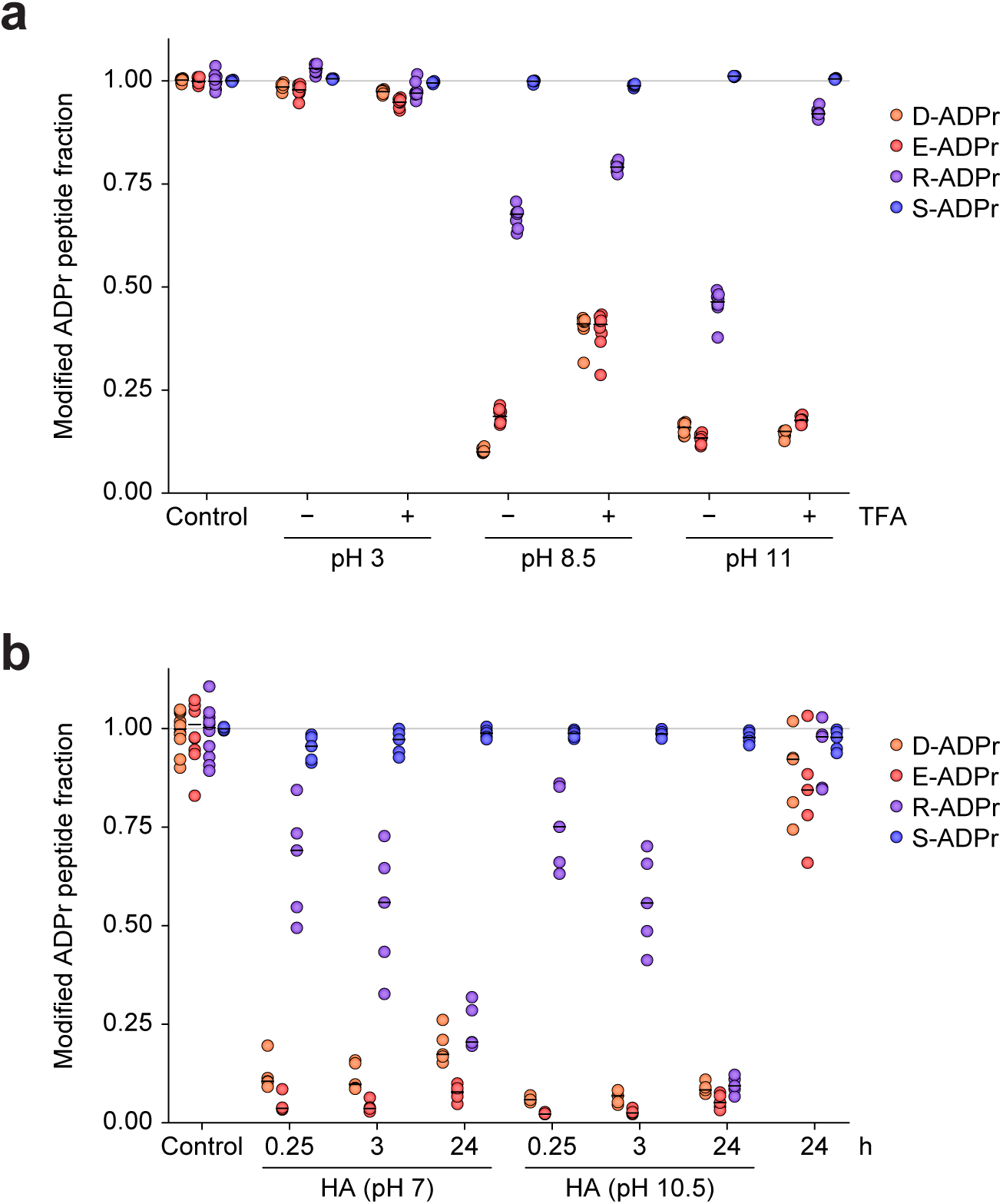
MS-based evaluation of stability of ADPr linked to Asp, Glu, Arg, and Ser residues. (A) Relative ADPr stability across different pH values across common MS buffers, vacuum dried in their respective buffers or otherwise first acidified using TFA. *n*=7 technical replicates. (B) Stability of different ADPr linkages following treatment with neutralized (pH 7) or standard hydroxylamine (pH 10.5), *n*=5 technical replicates. “HA”; hydroxylamine.

**Figure S2.**
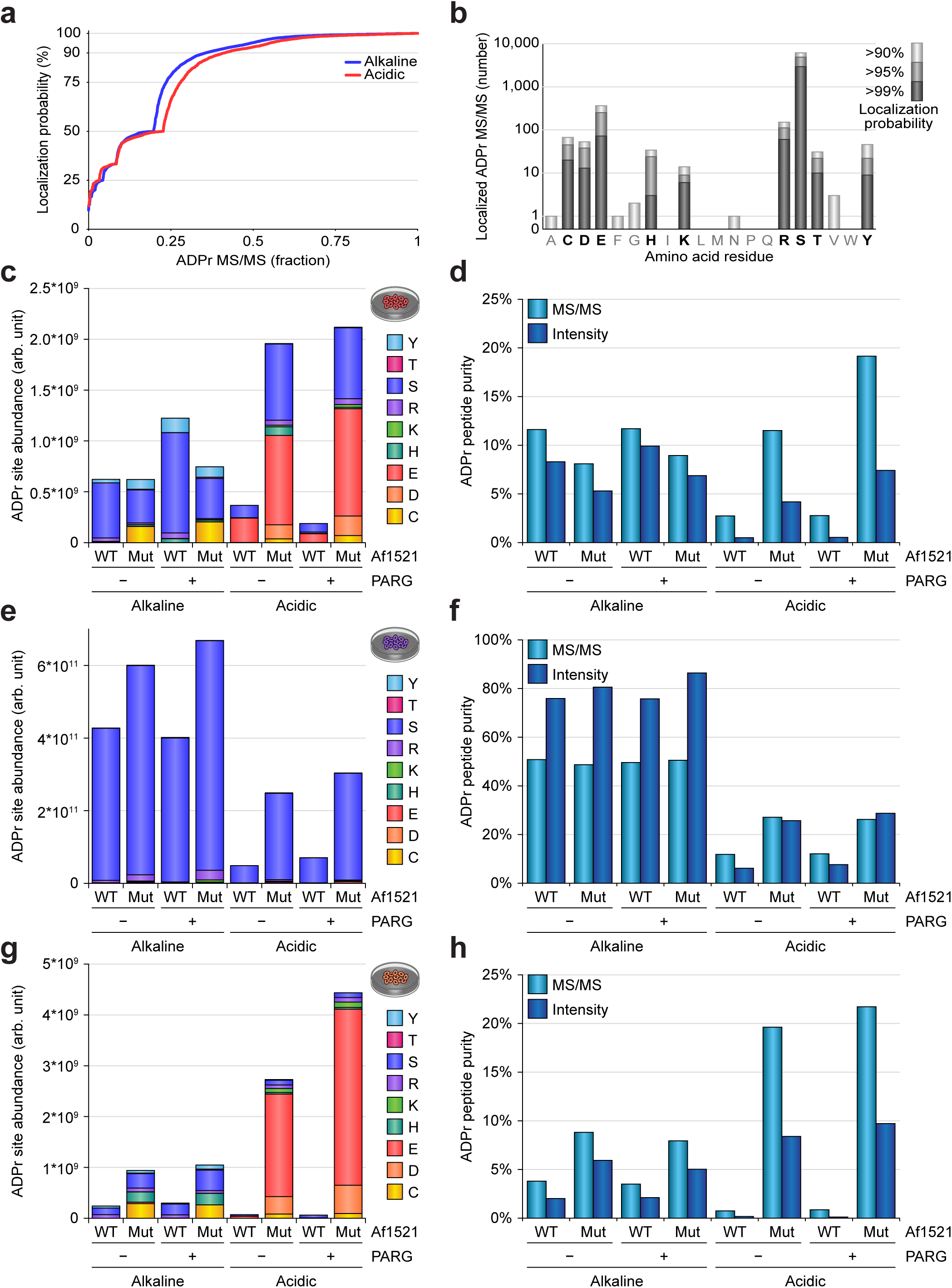
Benchmarking different proteomics workflows for purification of any ADPr linkage type. (A) Waterfall plot, visualizing the relation between all identified ADPr-modified MS/MS in a condition (x-axis), and the maximum localization probability of ADPr within each peptide (y-axis). (B) Number of ADPr-modified peptide-spectrum-matches (MS/MS) localized in a fully unrestricted search, wherein ADPr is allowed to reside on any residue type at any position in the peptides. For determining which amino acid residues are plausibly modified, we only accepted localization >99%. (C) Overview of the total ADPr site abundance (or “intensity”) profiled in the A549 +IFNγ samples, with distribution of amino acid types indicated. (D) Peptide purity analysis, plotting the MS/MS-based purity, derived by dividing the number of ADPr-modified MS/MS by the total number of MS/MS, and plotting the intensity-based purity, derived from dividing ADPr peptide intensity by the total intensity observed for the experiment. (E) As **C**, but for HeLa +H_2_O_2_ samples. (F) As **D**, but for HeLa +H_2_O_2_ samples. (G) As **C**, but for U2OS ΔHPF1 +H_2_O_2_ samples. (H) As **D**, but for U2OS ΔHPF1 +H_2_O_2_ samples.

**Figure S3.**
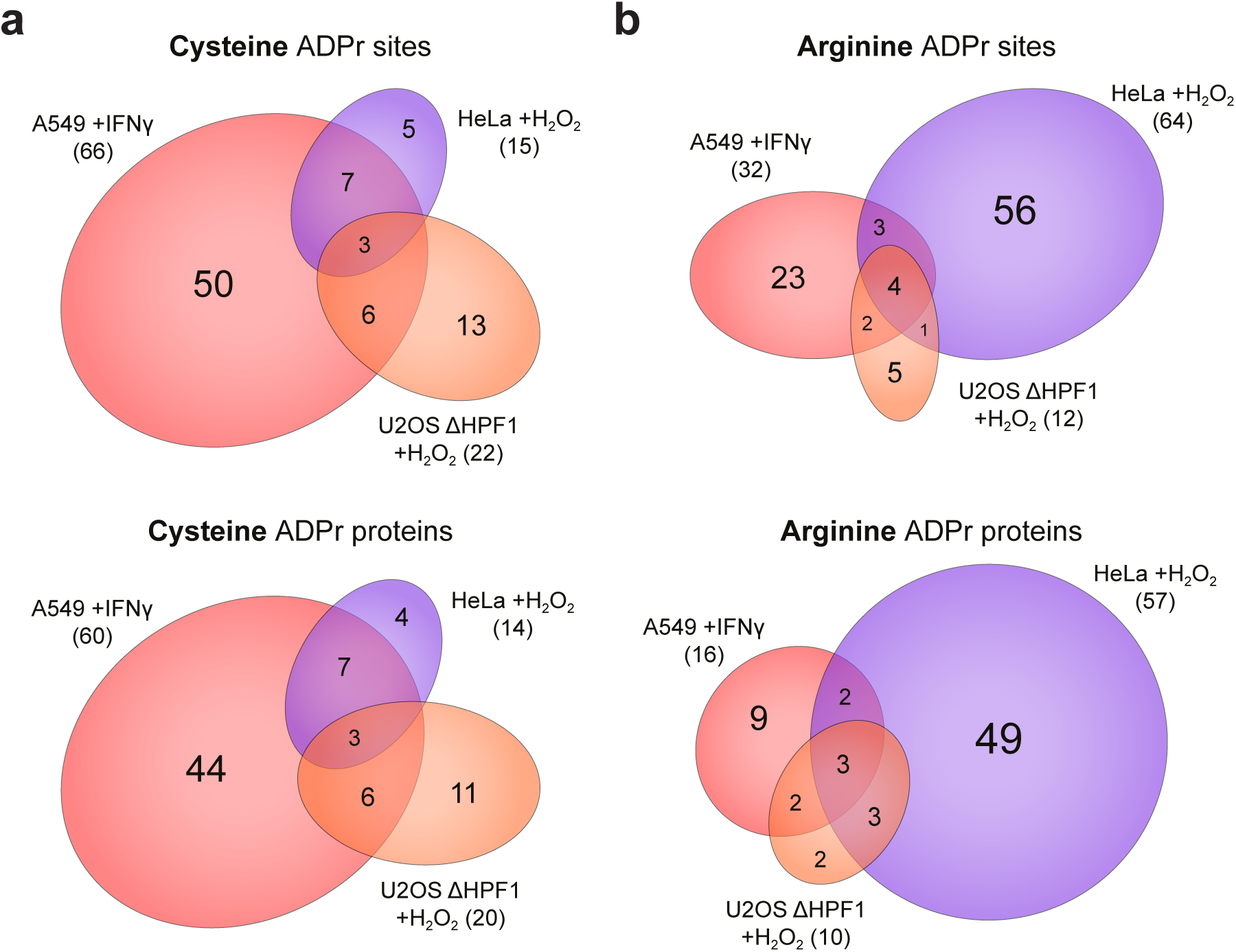
Distribution of different ADPr linkages across different cell lines and treatments. (A) Scaled Venn diagram showing overlap between the different cell types and treatments, for either Cys ADPr sites (top) or Cys ADPr target proteins (bottom). (B) As **A**, but for Arg ADPr.

**Figure S4.**
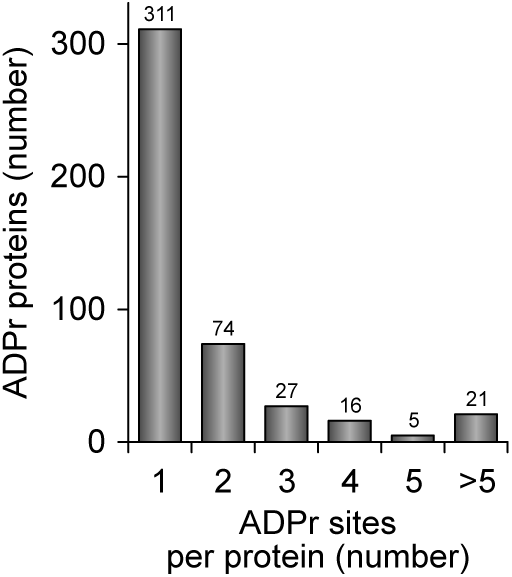
Overview of the number of ADPr sites identified per protein.

**Figure S5.**
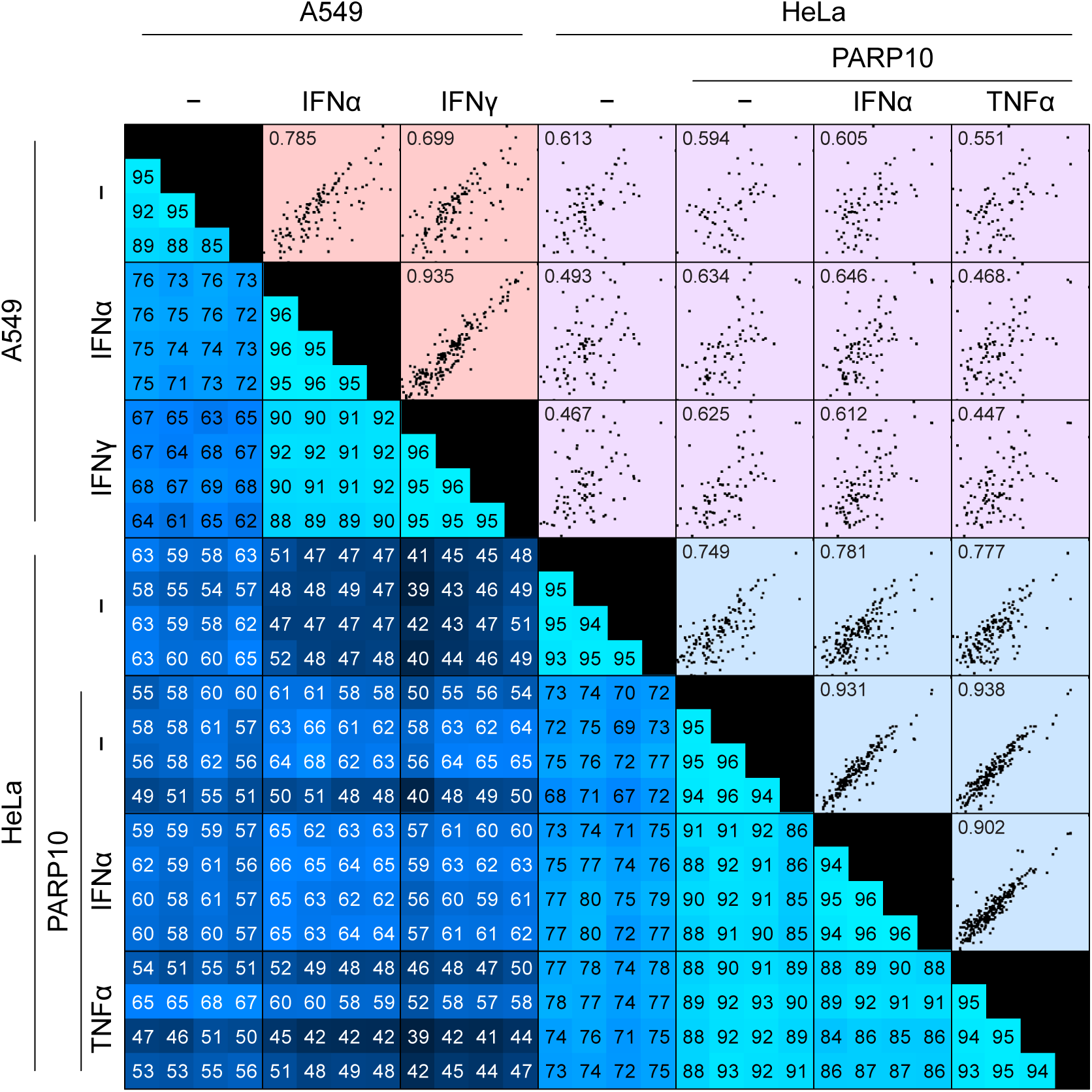
Global reproducibility analysis based on Pearson correlation. The bottom-left visualizes the degree of Pearson correlation (in %) between each replicate, across all conditions. Brighter blue corresponds to higher Pearson correlation. The upper-right shows a collection of scatter plot analyses comparing the abundance values of all ADPr target proteins between each combination of two experimental conditions. Within each scatter plot, protein abundances increase from bottom to top and from left to right, i.e. the most abundant proteins appear in the top-right corners. The number in each scatter plot indicates Pearson correlation as determined by linear regression.

**Figure S6.**
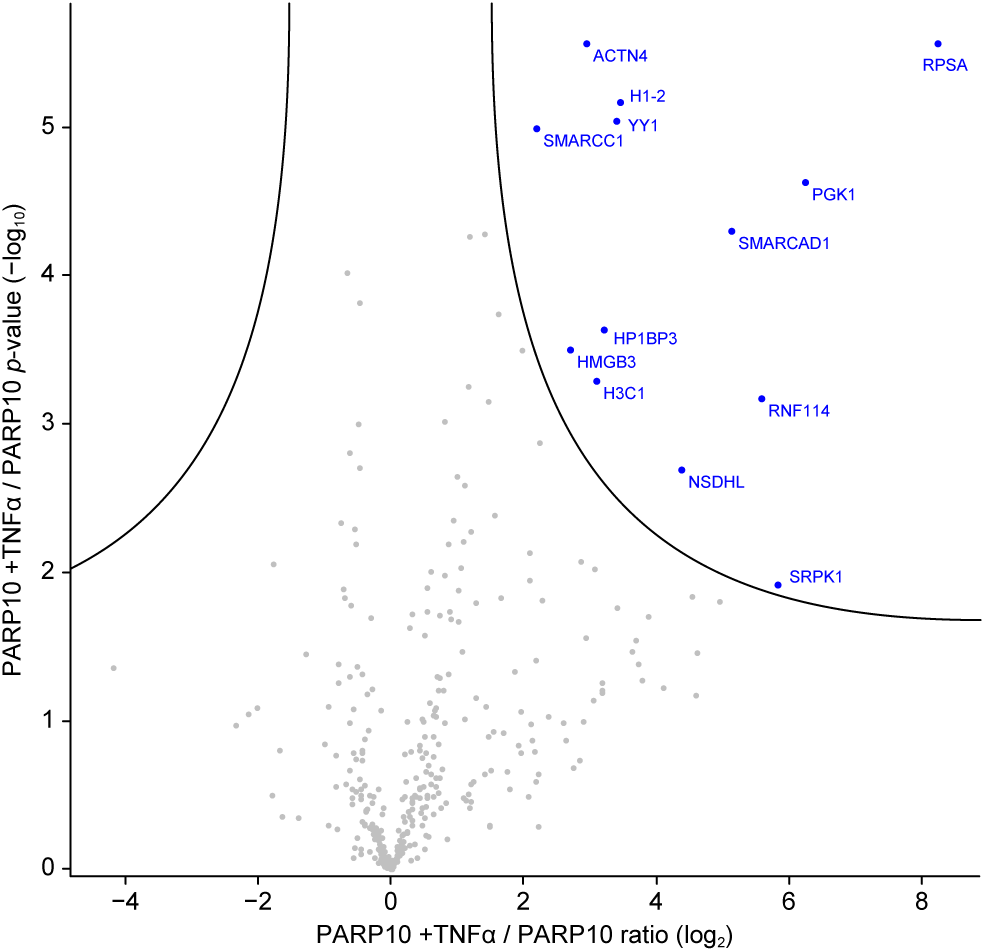
Volcano plot analysis, visualizing ADPr target protein dynamics between PARP10-induced +TNFα (right) and PARP10-induced (left). The significance cut-off line is drawn at *q*=0.01. Significance determined via student’s two-sided t-testing with permutation-based FDR control, *n*=4 cell culture replicates.

**Figure S7.**
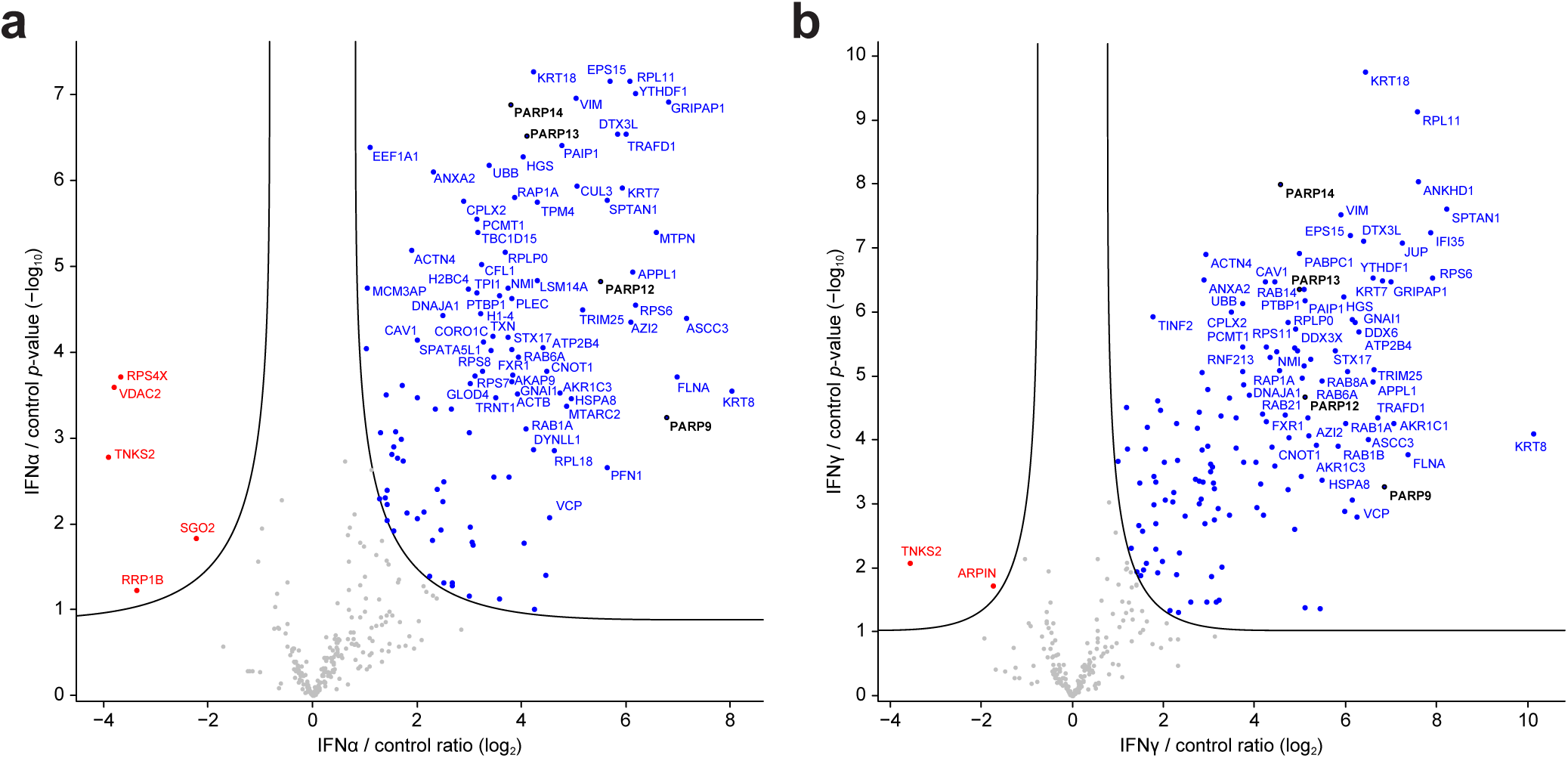
(A) Volcano plot analysis, visualizing ADPr target protein dynamics between A549 cells +IFNα (right) and control A549 cells (left). The significance cut-off line is drawn at *q*=0.01. Significance determined via student’s two-sided t-testing with permutation-based FDR control, *n*=4 cell culture replicates. (B) As **A**, but with IFNγ treatment.

**Figure S8.**
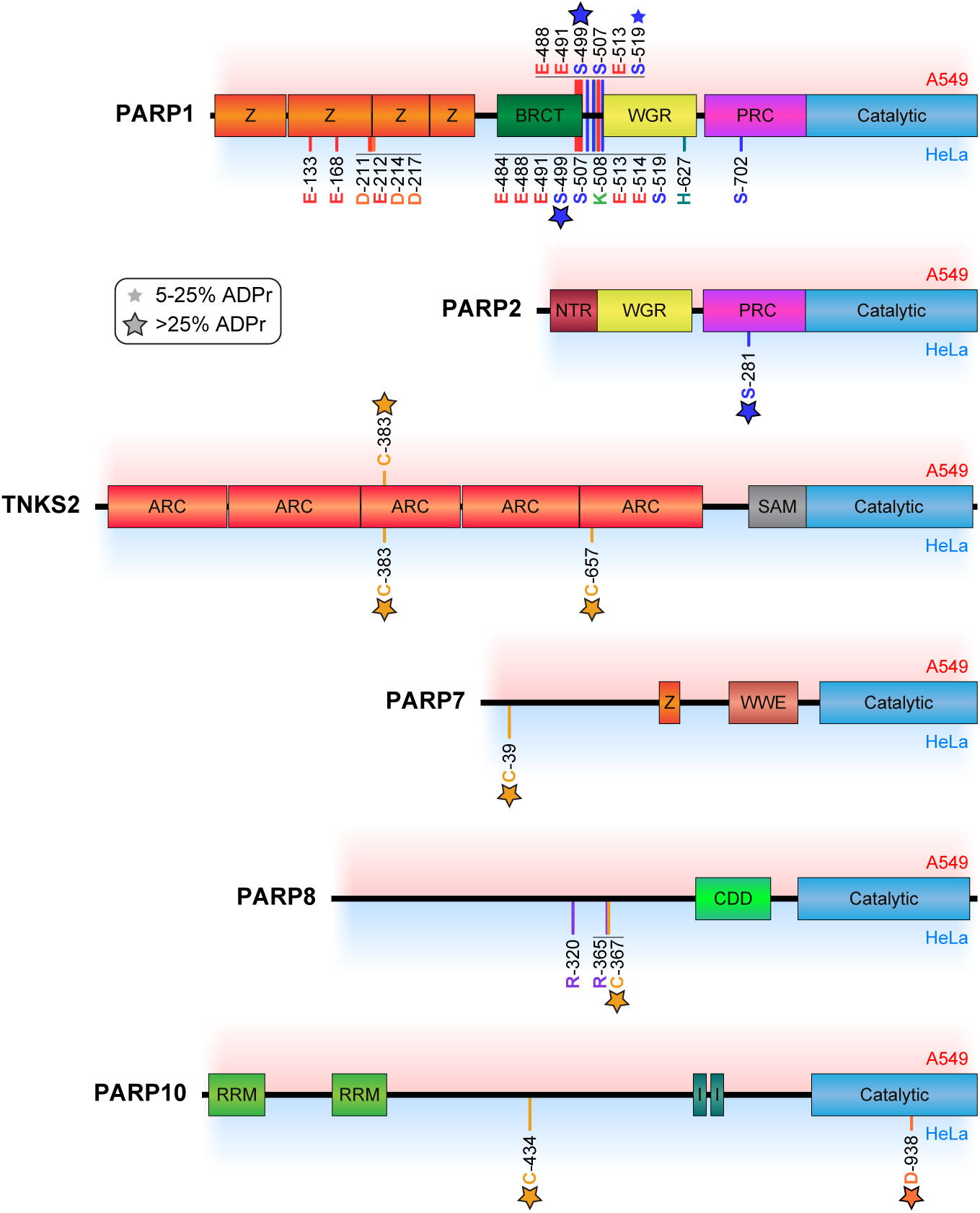
Linear visualization of PARP1, 2, 5B, 7, 8, and 10. Drawings are scaled to the length of the proteins, with known domains indicated. All mapped ADPr sites are marked, stratified by detection in A549 or HeLa. Small stars indicate 5-25% of total protein ADPr detected on the residue, whereas large stars indicate >25% of protein ADPr on the residue.

